# Formation, collective motion, and merging of macroscopic bacterial aggregates

**DOI:** 10.1101/2021.06.09.447728

**Authors:** George Courcoubetis, Manasi Gangan, Sean Lim, Xiaokan Guo, Stephan Haas, James Q. Boedicker

## Abstract

Chemotactic bacteria form emergent spatial patterns of variable cell density within cultures that are initially spatially uniform. These patterns are the result of chemical gradients that are created from the directed movement and metabolic activity of billions of cells. A recent study on pattern formation in wild bacterial isolates has revealed unique collective behaviors of the bacteria *Enterobacter cloacae*. As in other bacteria species, *Enterobacter cloacae* form macroscopic aggregates. Once formed, these bacterial clusters can migrate several millimeters, sometimes resulting in the merging of two or more clusters. To better understand these phenomena, we examine the formation and dynamics of thousands of bacterial clusters that form within a 22 cm square culture dish filled with soft agar over two days. At the macroscale, the aggregates display spatial order at short length scales, and the migration of cell clusters is superdiffusive, with a merging acceleration that is correlated with aggregate size. At the microscale, aggregates are composed of immotile cells surrounded by low density regions of motile cells. The collective movement of the aggregates is the result of an asymmetric flux of bacteria at the boundary. An agent based model is developed to examine how these phenomena are the result of both chemotactic movement and a change in motility at high cell density. These results identify and characterize a new mechanism for collective bacterial motility driven by a transient, density-dependent change in motility.

**Author summary:** Bacteria growing and swimming in soft agar often aggregate to form elaborate spatial patterns. Here we examine the patterns formed by the bacteria *Enterobacter cloacae*. An unusual behavior of this bacteria is the movement of cell clusters, millions of bacteria forming a tiny spot and moving together in the same direction. These spots sometimes run into each other and combine. By looking at the cells within these spots under a microscope, we find that cells within each spot stop swimming. The process of switching back and forth between swimming and not swimming causes the movement and fusion of the spots. A numerical simulation shows that the migration and merging of these spots can be expected if the cells swim towards regions of space with high concentrations of attractant molecules and stop swimming in locations crowded with many cells. This work identifies a novel process through which populations of bacteria cooperate and control the movement of large groups of cells.

## Introduction

In populations of chemotactic bacteria, coupling of the directed movement of individual cells in response to nutrients or chemical stimuli gives rise to spatio-temporal collective phenomena, including swarm bands and aggregates (1–3). These macroscopic structures are the result of an emergent pattern of bacterial cell density that forms due to the coordinated movement and metabolic activity of billions of bacterial cells in an initially uniform environment. Both swarm band and aggregate formation rely on chemotaxis (1–4). These collective phenomena can even be predicted with analytical mathematical considerations and recreated with detailed computational models (3,5–10). Decades of work has resulted in a detailed and predictive understanding of bacterial collective phenomena, based mostly on work with bacterial species *Escherichia coli, Salmonella typhimurium* and *Myxococcus xanthus* (1–3,11). It is unclear whether the collective properties within these two strains encompasses the full range of collective behaviors observed in all chemotactic bacteria, or if our understanding of these behaviors extends to other species of chemotactic bacteria.

A variety of cellular motility rules have been attributed to the emergence of larger macroscopic properties via the collective organization of high local densities of cells. A main driver of bacterial pattern formation is chemotaxis; individual cells utilize flagella to move in a combination of runs in a straight line, interrupted by tumbles to randomly change the direction of swimming (12,13). At the molecular level, the switch between the run state and the tumbling state allows the bacteria to navigate towards increasing chemoattractant gradients (14). For instance, many swarm bands of *Enterobacteriaceae* species are the result of cells migrating up concentration gradients of nutrients following local depletion (15). The collective responses can also depend on the cell density and shape. For example, the surface swarming of *Bacillus subtilis* exhibits different morphologies, depending on the aspect ratios and surface densities of the cells (16). Moreover, single cell properties such as adhesion, also direct the nature of collective phenomena. During the initial stages, the fruiting bodies of *Myxococcus xanthus* spontaneously assemble through the adhesion of cells, when two collide with each other (17). This diversity of mechanisms governing the movement of individual cells has given rise to multiple macroscopic dynamics.

Bacterial cells are internally driven motile agents, and thus belong to the category of active matter. Active matter is a branch of non-equilibrium physics that considers microscopic rules and emergent macroscopic phenomena of energy consuming motile agents (18). Driven inorganic matter also lies in the realm of active matter and has been used to probe the effect of individual properties of agents on the collective. For instance, self propelled colloidal particles form aggregates, with size that linearly increases with particle speed (19). Often living and non-living systems obey similar individual rules, for example swarming and swimming bacteria at high densities and shaken granular materials belong to the same active matter category of self-propelled rods (20). Local motility interactions, which induce a distance dependent velocity alignment of moving agents, as dictated by the Vicsek model, give rise to emergent collective motion in multicellular organisms (21). This density driven motility transition has been observed in the schooling of fish, flocking of birds, cells and insects (22). The concepts of active matter systems can help to understand complex biological and physical processes, and even help to develop micromachines and nanomachines for practical applications (23,24).

Here we report new collective properties observed in the bacterial strain *Enterobacter cloacae,* i.e., the formation, long-distance movement, and merging of macroscale bacterial aggregates. Chemotactic pattern formation was reported in this strain as part of a recent study of pattern formation in wild bacterial isolates (6). Here we quantify a pattern of spots that emerges within an initially well-mixed culture of cells in soft agar, and track the movement of each spot over multiple hours. In contrast to bacterial aggregates of *Escherichia coli,* which have been reported as mostly stationary with slight “jiggling” over time (1), aggregates formed by the strain *Enterobacter cloacae* migrate over distances up to four times their diameter. In addition, this movement results in approximately 36% of the aggregates merging with another aggregate to form a single spot. Our aim is to quantify and explain the spatial characteristics of these aggregates, their motility, and the underlying kinematics of the merging phenomena. High magnification, time-lapse imaging of individual cells within the spots reveal the microscopic mechanism that enables the collective motion of spots, namely a transition of individual cells between the motile and non-motile state. Chemotactic agent based simulations, which include a novel immotile to motile transition, recreate the spatial order observed in experiments. This transition gives rise to a novel type of collective motility in bacteria.

## Results

*Enterobacter cloacae* is a bacterial strain previously reported to be capable of large scale pattern formation, including moving bands of high cell density and aggregate formation (6,25). Here we focus on the aggregate formation, using a large culture dish and uniformly mixing cells into the culture medium at the beginning of the experiment. Cells grown overnight in Luria Bertani media at 37°C and 180 rpm were uniformly mixed into soft minimal agar media supplemented with glucose at 1.5% inoculum. The media containing cells were poured into 22 cm × 22 cm dishes, with a lid and incubated at room temperature, as depicted in Fig. 1A. Pictures of the dish were taken over time using a DSLR camera at 1X magnification. Initially, no macroscopic aggregates could be observed. After 5 hours, aggregates began to form, and at around 20 h a pattern of spots emerged across the plate, as shown in Fig. 1B and C. Images of aggregates were analyzed using image analysis software as discussed in methods.

**Figure 1:**
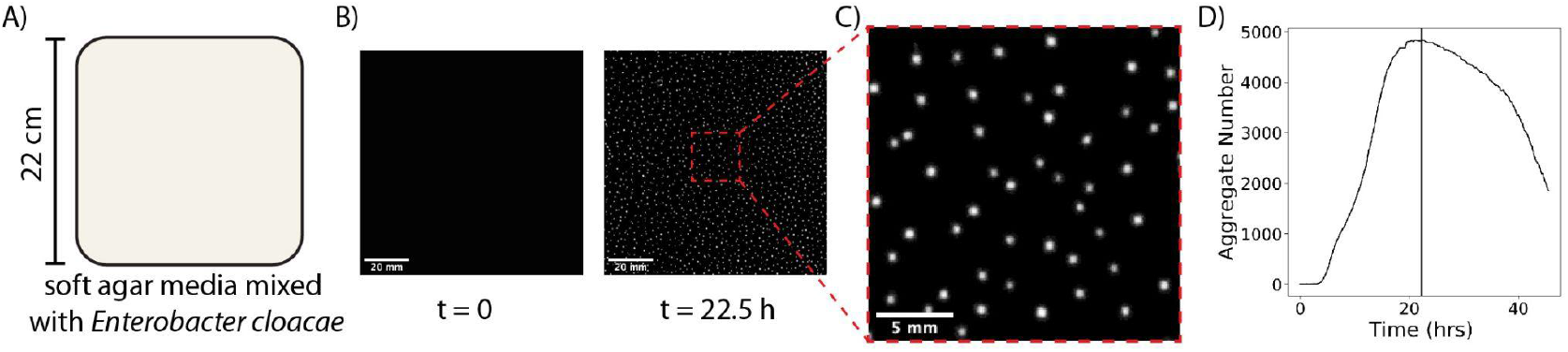
Overview of aggregate formation experiment. **A)** Sketch of experimental setup. **B)** Image of a 100×100*mm*^2^ subregion of the experimental plate before and after aggregate formation. **C)** Zoomed in 20×20 mm^2^ region showing the aggregates at 22.5 h. **D)** Number of aggregates on the plate versus time for the duration of the 44 hour experiment. The number of spot aggregates peaks at 22.5 hours, marked by the vertical line.

Aggregates form over about 10 hours, reaching a maximal number of aggregates at 22.5 h, as shown in Fig. 1D. A video of spot formation over the entire plate can be found in SI Video 1. As shown in the video, many of the aggregates migrate on the plate after formation, and some even merge together. After 36 h, the aggregates begin to dissolve and are no longer visible on the plate. In the next sections, we will analyze the formation and the spatial patterns of these aggregates, the migration of aggregates on the plate, as well as the observed aggregate merging process.

### Aggregate spatial structure analysis

As shown in Fig 1, thousands of aggregates appear on the plate. We now analyze the distributions of sizes and nearest neighbor distances of the aggregates at the time of the maximum number of aggregates. However, as shown in SI Fig. 2, aggregate sizes and the overall spatial distribution did not vary significantly between the time when spots started filling the plate and when the spots began to dissolve. As observed in SI Fig. 3, there was a large scale variation in the local density of aggregates, the densest regions containing 25 aggregates per *cm*^2^ and the sparsest regions having 5 aggregates per *cm*^2^. Despite this variation in the density of aggregates, the trends observed in analysis of the center of the plate, shown in Fig. 2, are similar to trends in the larger plate (SI Fig. 2). The subset of the plate, highlighted in SI Fig. 3, was used to obtain a higher resolution of the local spatial structure and to avoid edge effects. As shown in Fig. 2A, the aggregates were fairly uniform in size, with an average diameter near 1 mm. A small fraction of aggregates is larger and does not belong to the same peak in the histogram. Nearest neighbor distances were calculated using Voronoi tessellation, considering all nearest neighbors (26). As shown in Fig. 2B, the distribution of nearest neighbors is a slightly skewed Gaussian, with a characteristic nearest neighbor distance around 2.7 mm.

**Figure 2:**
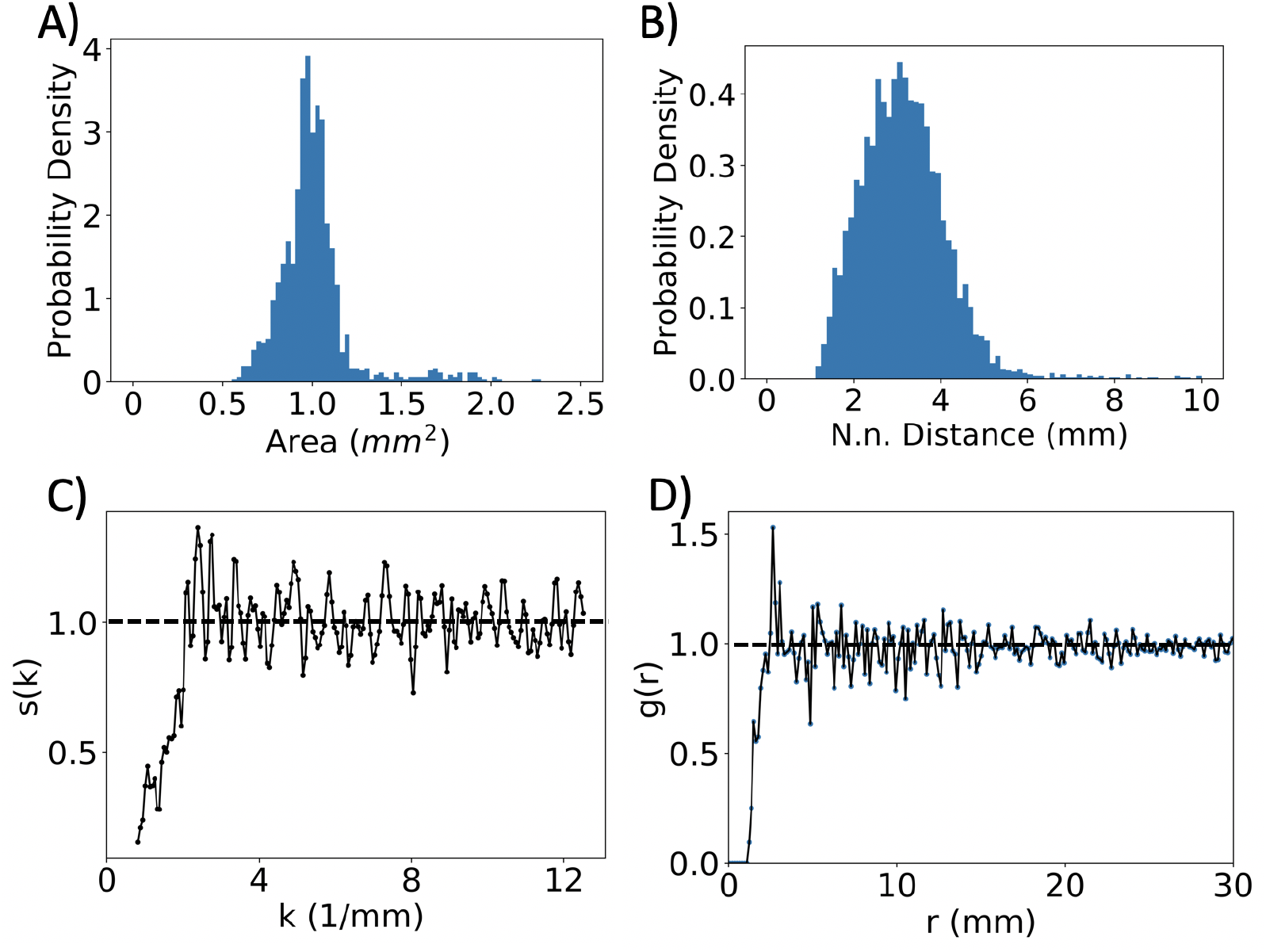
Spatial structure of aggregate formation. **A-D)** Spatial analysis of the aggregate pattern at 22.5 h for a subregion of the plate. **A)** The aggregate area distribution is depicted, with a mean of 1.01 *mm*^2^ and standard deviation of 0.25 *mm*^2^. **B**) Histogram of the nearest neighbor distance distribution. A pronounced single peak shows the existence of small scale structure in the pattern. The average is 3.33 *mm*, with standard deviation of 2.33 *mm*. **C)** Structure factor versus wavenumber. The structure factor saturates to 1 at 3 *mm*^-1^, suggesting absence of long range order. **D)** Radial pair correlation function. The pair correlation is zero for distances less than 1.7 average aggregate diameters, as no other spots are detected in that proximity. The pair correlation peaks at 3.37 mm, exhibiting short range order, and saturates to one at 7 *mm*, consistent with disorder at longer distances.

To further analyze the overall spatial pattern of aggregates, the pair correlation function and the structure factor of the subregion of the plate were calculated (see methods for details). As observed in Fig. 2C, the structure factor shows a single peak at 2. 5 *mm*^-1^, and lacks additional peaks, indicating an absence of long range order. This type of structure factor resembles that of a liquid, characterized by a single peak at low wavenumber, k, that repeats with a diminishing amplitude for multiples of nearest neighbor distances. However, it is unclear whether the fluctuations for large k are due to noise and finite sample size or long range order. Fig. 2D shows the results of the radial correlation analysis. Here, the ultra-short aggregate-to-aggregate distances appear to be absent, indicating an exclusion zone approximately two times larger than the average aggregate diameter. The short range structure identified in the pattern matches that of a liquid, which is described by an exclusion region mediated by short range repulsion. In the long range limit, the pattern of aggregates resembles a gas, as there is no long range order. In summary, the aggregate pattern can be described as having no Bragg peaks, i.e., no long range order (Fig. 2C), but instead short range correlations and a restriction on the minimal spacing between aggregates.

Hard sphere models have been widely used to model liquids and successfully capture their quasi-universal spatial structure (27). In the hard sphere model, each particle is defined as a sphere with a fixed radius, which cannot overlap with the other spheres in the system. Interestingly, the spatial structure of the bacterial aggregates is consistent with a system of closed packed hard spheres whose radii are distributed according to a Gaussian (28). Specifically, the coefficient of variation for spot size distribution was determined to be η = 0. 3, and using the same coefficient of variation for a Gaussian, size distributed hard sphere packing, one retrieves an identical functional form for the radial pair correlation function (28). This is consistent with the experimental aggregates size distribution. However, contrary to the hard-sphere model, the bacterial aggregates are not closely packed, and nearest neighbors are on average separated by approximately three times the average aggregate diameter. The explanation for this difference is that the aggregates form by recruiting bacteria in proximity, thus extending their exclusion region beyond the physical aggregate size.

### Aggregate motility analysis

After aggregate formation, a displacement of some of the spot aggregates over time was observed, as shown in Fig. 3. In Fig. 3A, the green circle shows the location of each aggregate at 14.5 h, the magenta circle shows the location of each aggregate at 37.3 h, and the yellow line indicates the aggregate trajectory. Over twenty hours, the aggregates were displaced by up to a few millimeters, without a significant change in aggregate area or shape. Note that the upper left aggregate dissolves prior to 37.3h.

**Fig 3.**
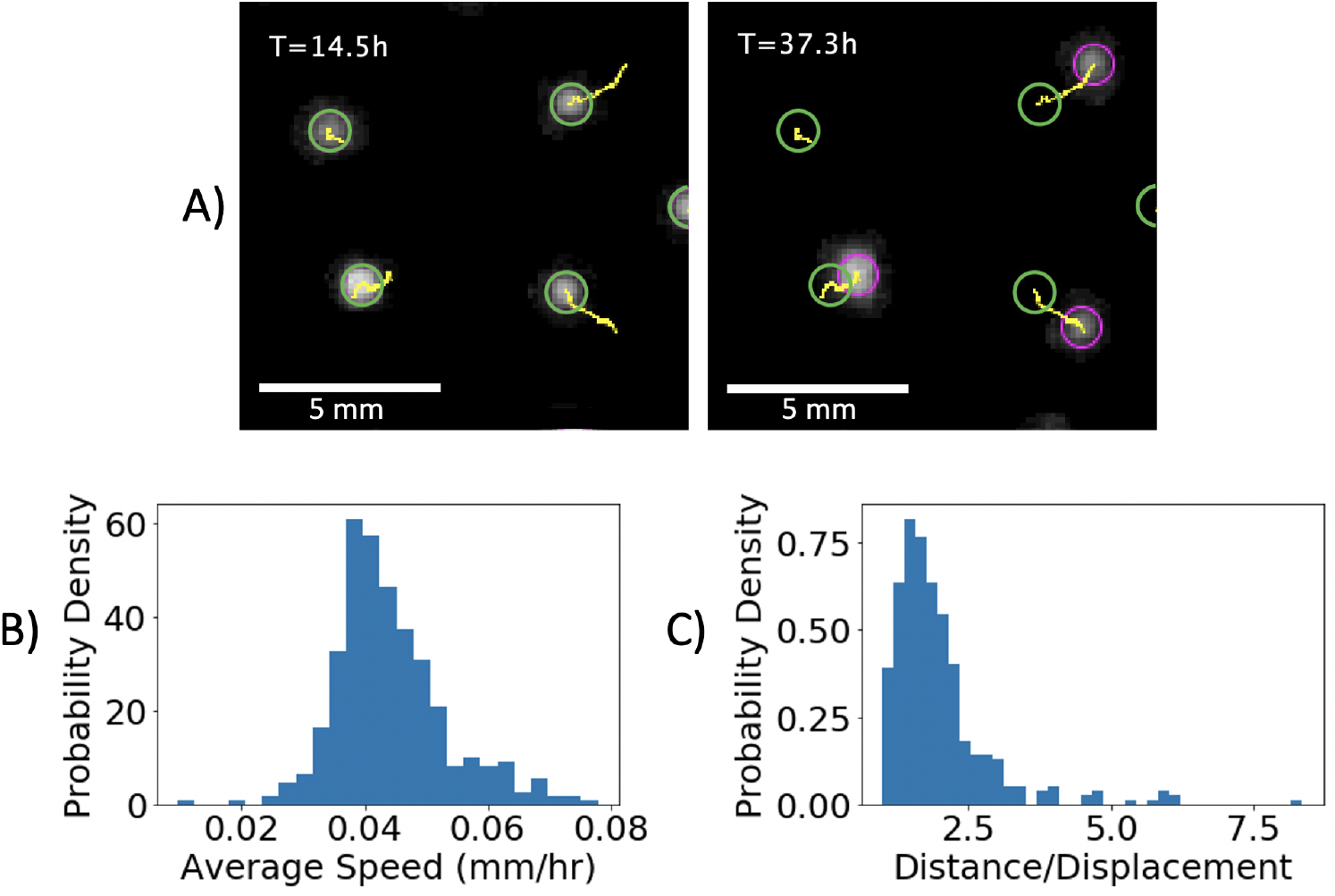
Aggregate motility. **A)** Snapshots of a subregion of the plate at T=14.5 h and T=37.3 h. The aggregates are depicted along with the trajectory identified by the tracking algorithm. **B)** Distribution of the average speed of all aggregates. **C)** Distribution of path deflection, quantified as the ratio of distance to displacement. The path taken by the aggregates slightly deviates from a straight line.

Time-lapse movies of the dish were used to quantify the trajectories of 495 aggregates, obtained by tracking the aggregates in six 26 mm × 26 mm subregions, shown in SI Fig. 4. An analysis of the aggregate trajectories reveals that a subset of 35.4% of the aggregates coalesced or merged. In the next sections we discuss the merging of two or more aggregates, while here we focus on the motility of all aggregates. Fig. 3B shows the distribution of speeds for all the aggregates. The average speed of aggregate migration is 0.044 mm/hr. The path taken by the aggregates is quantified by dividing the distance over displacement in Fig. 3C. Here, a value of 1, the minimal value, indicates a straight line trajectory. The ratio of distance to displacement shown here, with an average value of 2.0, indicates that aggregates move in a directed manner. Plotting the average mean squared displacement versus time yields that the aggregate trajectories lie in the super-diffusive regime with a power law coefficient of 1.5 (SI Fig. 5A). Isolating the non-mergers, a weak but statistically significant negative correlation is found between the average minimum nearest neighbor distance and average speed for each aggregate trajectory, with a spearman rank-order correlation coefficient correlation −0.12, p-value 0.034 (SI Fig. 5B). The negative correlation indicates that proximity with other aggregates increases the speed of a given aggregate.

To shed more light on the microscopic behavior of the cells within as well as around the aggregates, we scaled down the experimental system using the lab-Tek chamber (SI Fig. 6) and observed the movement of over 40 aggregates at high magnification. For this experiment, cultures were prepared by tagging 5% of the populations with RFP, so that single cell movements can easily be monitored (29). The presence of some fluorescent cells also enables the detection of the aggregate boundary, as high density regions of cells have a higher fluorescent intensity caused by cells outside of the focal plane. Note that the growth rate as well as average speed of *E. cloacae* cells transformed with RFP expressing plasmid is comparable to that of wild type *E. cloacae* cells (SI Fig. 7) ensuring the reproducibility of the experiment in the lab-Tek chamber with mixed populations.

Aggregates were imaged and their displacements were analyzed over 80 mins (Fig. 4A, SI Videos 3 and 4). ImageJ was used to determine the boundary of a spot at different time frames. Then, the difference between the xy-coordinates of the centroid of the boundary in the initial frame (solid cyan blue line) and in the final time frame (solid yellow line) were calculated to obtain the boundary propagation speed. An analysis of 40 aggregates reveals that the average speed exhibited by the aggregate front is 0.023 mm/ hr (Fig. 4D), which is comparable to the aggregate speed calculated for large plate experiments. Since the microscope was focused at a random segment of the aggregate boundaries, there is no guarantee that the trailing or receding edge of the aggregate was recorded. Therefore, measuring the boundary propagation speed corresponds to obtaining a component and not the magnitude of aggregate speed.

**Figure 4:**
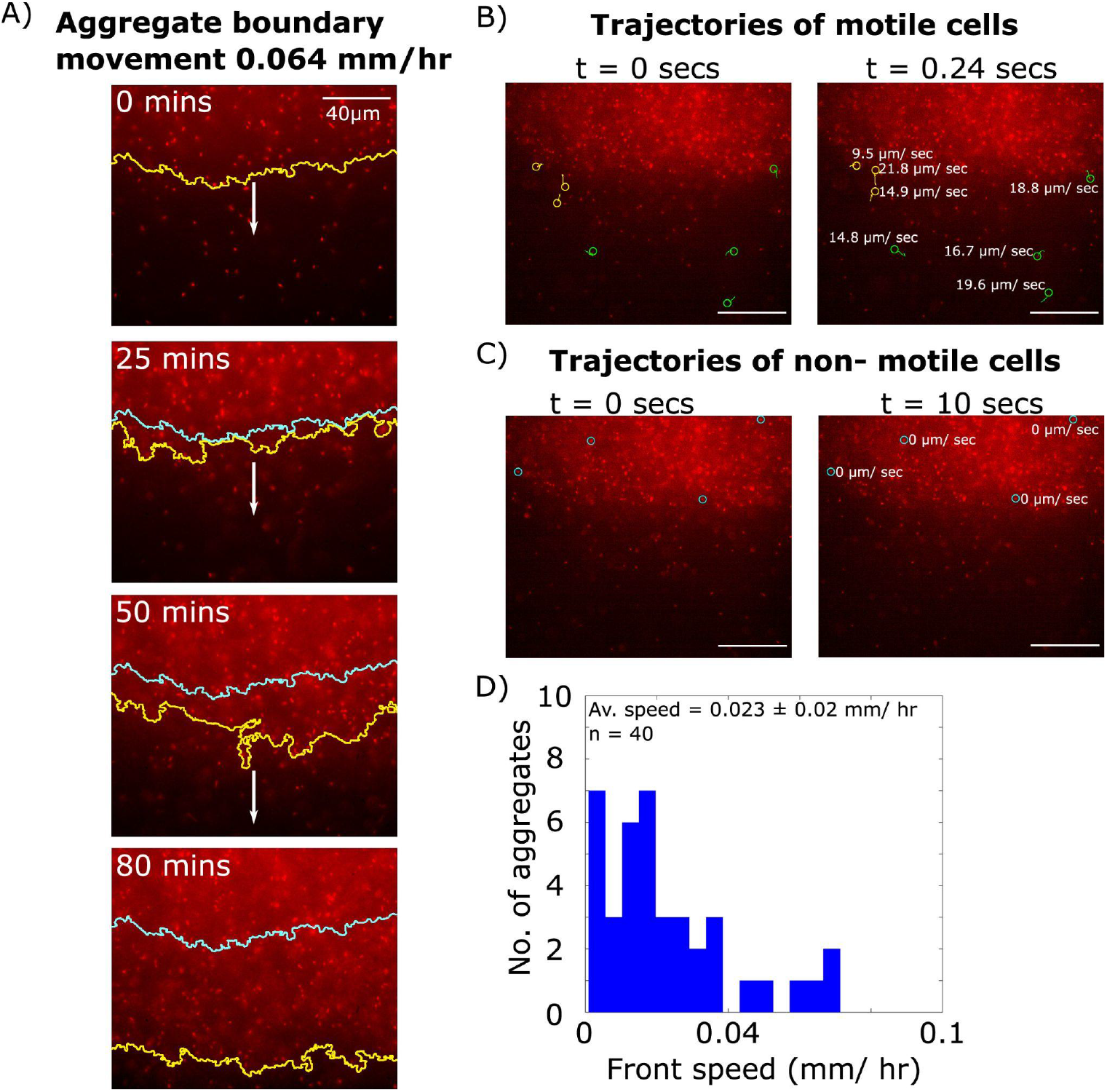
Single cell motility and aggregate movement. High magnification imaging of cells in an aggregate. Aggregates were formed within a coverslip bottom chamber to enable 100X imaging of cells within the aggregate. (A) Aggregate movement on the microscale. Progression of the boundary of a typical aggregate over 80 mins. The yellow solid line marks the periphery of the spot in real time, whereas the cyan solid line indicates the position of the spot front at t = 0 mins. Direction of the spot movement is shown with the white arrow. (B) Motile cells exist in the vicinity of the aggregate and a subset consolidate with the aggregate and become immotile. Representative trajectories of individual cells that are motile at t=0. Yellow lines denote cells that swim towards the spot and become immotile within 0.24 s. Green lines denote cells that remain motile for 0.24 s. (C) Immotile cells within the aggregate. Shown in blue are representative cells within the aggregate that do not change position over 10 seconds. In B and C, trajectories are labeled with respective cellular velocities in μm/sec, and the scale bar is 40 μm. (D) Distribution of aggregate front speeds. By tracking the position of an aggregate boundary over time, the front speed is calculated for n=40 aggregates. The histogram shows the measured distribution of front speeds with an average value of 0.023 mm/hr.

Analysis of time-resolved videos help us track individual cells through time. Displacements calculated for single cells reveal the existence of three sub-populations in the milieu (Fig. 4B and C). As shown in Fig. 4B, a typical spot was observed to be surrounded by motile cells (a subset of such cells identified with green circles). Interestingly, some of the motile cells coalesce with aggregate and lose their motility, thus switching into non-motile cells (yellow circles). As shown in Fig. 4C, cells within the boundary of the aggregate were found to be immotile for the duration of the video (cyan blue circle). These microscopic observations suggest spot movement must be driven by the flux single bacterial cells leaving and joining the aggregate. In the direction of movement, individual cells join the aggregate and lose motility (SI Video 3), whereas at the trailing edge motility is regained and cells leave the aggregate (SI Video 4).

### Aggregate merging analysis

Next, we analyze the merging of two or more aggregates. The subregions studied for quantifying spot motility were also used for analyzing merging events (SI Fig. 4). As shown in Fig. 5A, the movement of aggregates sometimes results in the combination and merging of multiple aggregates into a single aggregate. Fig. 5A shows merging of two sets of aggregates. Upon analysis of the trajectories for 495 aggregates, 64.6% of aggregates did not merge, 31.1% aggregates merged with another aggregate, and 4.2% of aggregates merged with two other aggregates. Examples of mergers involving more than two aggregates are shown in SI Fig. 8. The remainder of the analysis is done for merging events with one other aggregate, which we define as two-spot mergers. Within this subset, only 46 out of the 77 trajectories were included in the analysis, i.e., leaving out merging trajectories with less than 20 positional data points whose dynamics could not be determined with a similarly high degree of accuracy.

**Figure 5.**
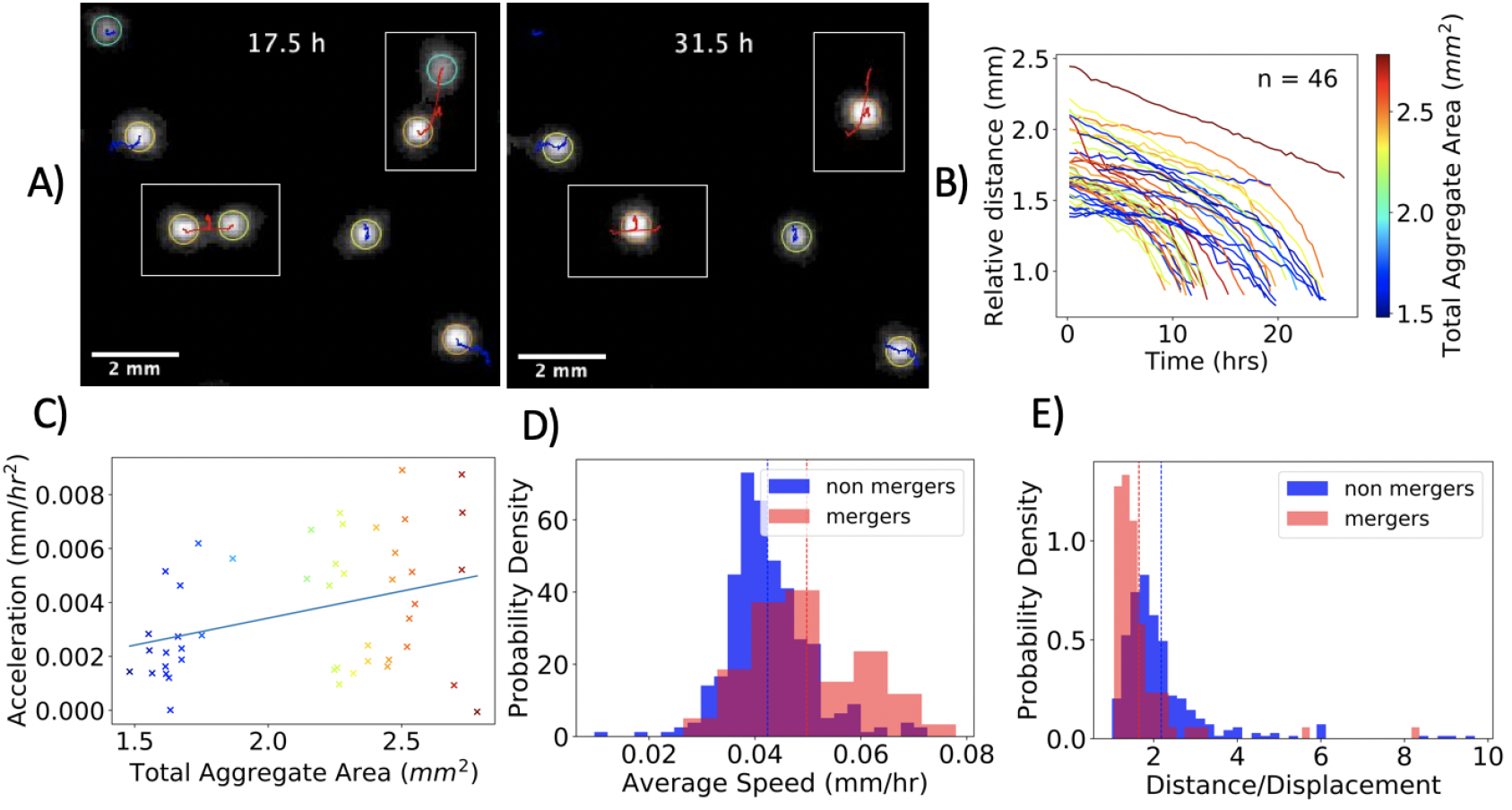
Aggregate Merging. **A)** Snapshots of a 10 mm × 9mm subregion of the plates at T=17.5 h and T=31.5 h. Two instances of two-aggregate merging events are highlighted in the boxed regions. The trajectories are plotted in blue for non-merging aggregates and in red for mergers of two aggregates. **B)** Relative distance vs. time for all merging processes, color coded with total aggregate area. **C)** The acceleration is positively correlated with the total aggregate area (spearman’s rho 0.30 and p-value 0.040). **D)** Distribution of aggregate speed for two-spot mergers and non-merging spots. The average (shown with a dashed line) illustrates that aggregates that eventually merge, on average, move faster. **E)** Distribution of trajectory distance divided by displacement for two-spot mergers and non-merging spots. On average, the path taken by aggregates that eventually merge is more direct.

The trajectories taken by merging aggregates are shown in Fig. 5B. The relative distance of the two merging aggregates is defined as 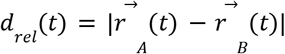, where 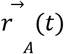 and 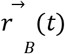 are the two-dimensional position vectors of aggregate A and B at time t. The trajectory taken by most merging aggregates was similar, exhibiting an increase in aggregate velocity as the distance between aggregates decreased. To quantify the dynamics of the merging spots, it was approximately treated with a constant acceleration model. Thus, the relative distance versus time for each trajectory was fit to a quadratic function. The quadratic fit was excellent for the majority of trajectories, with an average error of 0.15 pixels. The fit revealed the acceleration of each aggregate, and as shown in Fig. 5C, the acceleration was found to be correlated with aggregate size (Spearman rank-order correlation coefficient correlation 0.3, p-value 0.040). Quantifying the relative position of the aggregates relative to point of collision gave similar results, see SI Fig. 9. Curiously, unlike inertia dominated systems, the smaller spots do not necessarily move more than larger spots (SI Fig. 10). As shown in Figs. 5D and 5E, the aggregates that merged moved fast on average and took a more direct path. The non-merging aggregates had a mean speed of 0.042 mm/hr, whereas the mergers had a mean speed of 0.050 mm/hr, with a statistically significant difference of 0.007 mm/hr (p=0.0001 for an unpaired t-test). The fact that merging aggregates are on average faster and follow a more direct path suggests the presence of an effective attractive force between aggregates.

### Simulation

Based on the experimental observations, a 2D agent-based model was developed to explain the phenomena observed in aggregates of chemotactic bacteria. Our simulations were based on a model proposed in (9), with algorithms discussed in (30–32). In the simulations, 100,000 agents (cells) perform random brownian motion while experiencing a chemotactic drift force. Regarding chemotactic sensitivity, receptor law sensitivity was used (9), i.e., the chemotactic force magnitude increases with the slope of the chemoattractant gradient but decreases with the concentration, a formulation motivated by the saturation of bacteria surface receptors. Chemoattractant is produced and expelled by each agent, and the chemoattractants undergo diffusion and decay over time. The model does not incorporate cell division, death or food gradients. Further rules were implemented to make the simulation more realistic and capture experimental results: agents were attributed a finite size, they interact via a hookian repulsive force to simulate finite size, and they experience a chemoattractant dependent motility transition (See Appendix 1 for detailed model description).

Based on our observation that cells within the aggregates are immotile, the model was generalized to include a mechanism whereby an agent can transition from being motile to immotile and visa versa. To that end, a chemoattractant threshold was introduced, beyond which bacteria transition to an inmotile state. Since the chemoattractant concentration is proportional to the local density inside the aggregates, using a chemoattractant threshold corresponds to a density-dependent motility transition. As shown in SI Fig. 11, introduction of the density-dependent motility transition resulted in a greater fraction of motile cells within the population. Including only the density-dependent transition to the immotile state would lead to the eventual absorption of the vast majority of cells in aggregates. In experiments, however, the aggregates coexist with freely swimming cells throughout the course of the experiment. Furthermore, the analysis of aggregate movement in Fig. 4 indicates examples of the departure of cells from the aggregate interface. Therefore, a rule for reactivating motility was needed, even within high cell density regions. There are examples in the literature of a timed motility switch (33), so a rule was incorporated for motility to be regained after a random interval of time. More specifically, an agent stays in the immotile state for a fixed interval of time, after which there is a constant probability of regaining motility per iteration. To enable newly mobilized cells to leave regions of high cell density after regaining motility, cells ignore the chemoattractant gradient for a short period, performing brownian motion, after switching from the immotile to motile state. The expected value for the time scale for an agent to regain motility was set to 33 mins and the time scale for random walk after regaining motility was set to 15 mins.

The spatial scale of the simulation was set by equalizing the average aggregate radius of the simulation with the experiment. The time scale was set by equalizing the average speed of an agent with that of *Enterobacter cloacae,* as measured in (12). To perform this scale calibration, the average speed of every motile agent was calculated from the simulation, and equated to the experimental value. The parameters were chosen by numerical testing and are shown along with a laconic description in SI Table 1. The goal of the numerical exploration was to retrieve formation of aggregates while qualitatively capturing the relative aggregate size and spacing seen in the experiment. Exploration of systematically varying different parameters and components of the model are shown in SI Fig. 12–14. The model has 13 parameters and is non-linear and stochastic so it is simply not feasible computationally to systematically test the entire parameter space and analyze the emergent properties. The goal of these simulations was not to extract and use precise numerical values for unknown parameters, but instead to explore whether an established model of chemotactic behavior combined with the experimentally observed motility transition would be able to qualitatively reproduce the experimentally observed phenomena.

Our simulations, using these rules for motility switching, reproduce the formation of bacteria aggregates observed in experiments. Over time, high cell density aggregates composed of mainly immotile cells form. After the formation of aggregates and throughout the simulation, motile and immotile cell populations coexist with approximately 40% of cells retaining their motility at any given time (SI Fig. 15). As a consequence of setting the spatial scale using the experiment, these aggregates have a diameter of around 1 mm. The size distribution resembles experimental results, but exhibits higher variance. As shown in Figs. 6B-C, the spatial order of aggregates observed in the simulations is similar to the experimental measurements. The pattern retains short range order, described by an exclusion region, resembling a liquid, and disorder at long spatial distances, resembling a gas.

**Fig 6.**
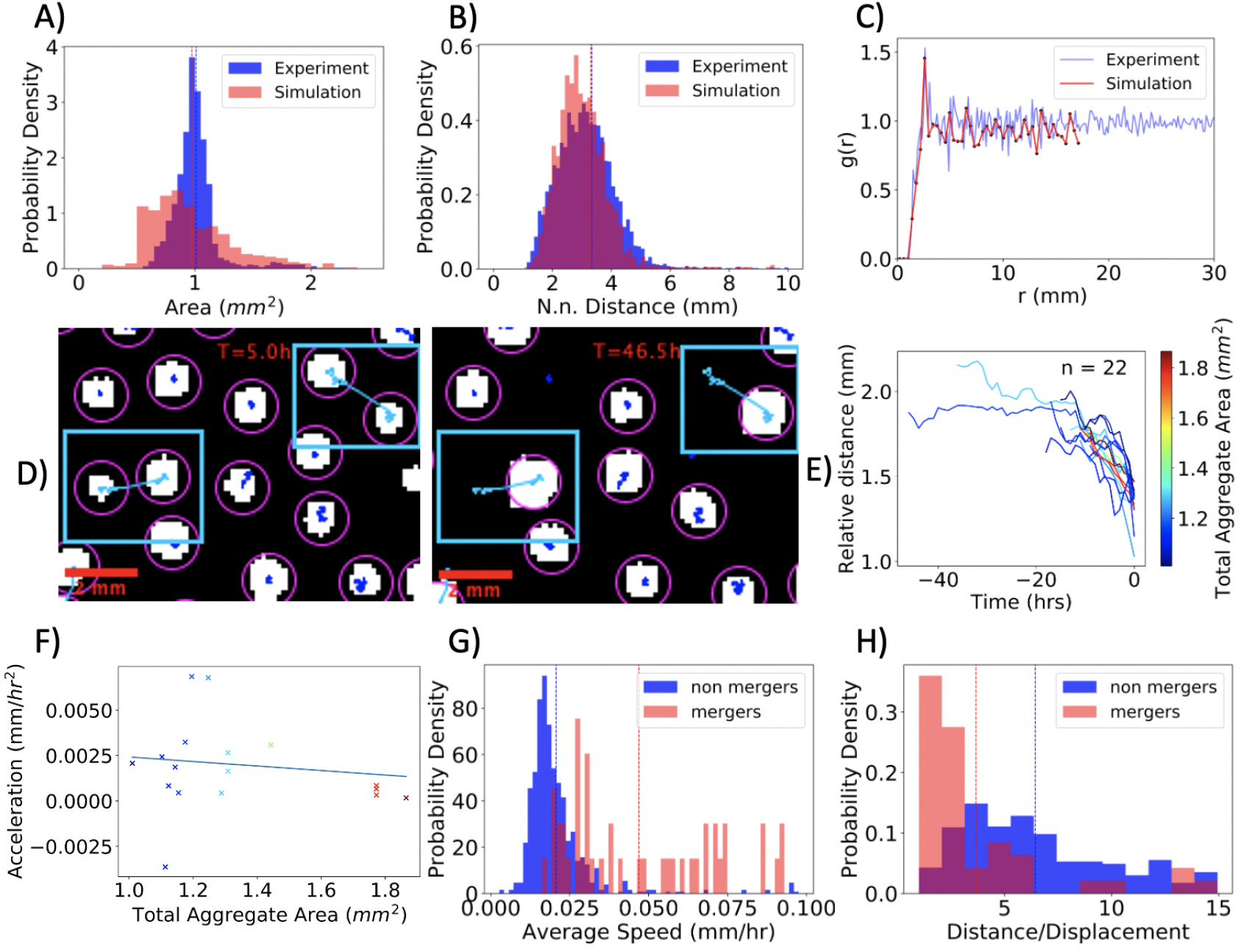
Computational Model: Spatial Structure and Merging. **A)-D)** Spatial Structure of Aggregates. **A)** The simulated spot area distribution has an average of 0.97*mm*^2^ and a standard deviation of 0.38 *mm*^2^. **B)** The simulated nearest neighbor distribution has an average of 3.30*mm* and a standard deviation of 2.49*mm*. **C)** The spatial correlation function of the simulated aggregates mirrors the saturation of the experimental structure, with saturation to 1 at 3.5mm. **D)** Snapshots before (T=5.0 h) and after (T=46.5 h) the merging of two segmented aggregates (see methods for segmentation details). **E)-H)** Merging dynamics in the simulations. **E)** Relative distance vs. time for all merging processes, color coded with total aggregate area. **F)** The acceleration is not found to be correlated with aggregate size (spearman’s rho 0.01 and a p-value of 0.962). **G)** Distribution of aggregate speed for two-spot mergers and non-merging spots. The average (shown by a dashed line) illustrates that mergers, on average, move faster. The speeds of the simulated aggregates are on the same order of magnitude as found in the experiments. **H)** Distribution of the ratio of trajectory distance divided by displacement for two-spot mergers and non-merging spots. On average, the path taken by spots that merge are more direct. Non-merging spots follow a less direct path as compared to the experiments.

In addition to forming a large-scale pattern of aggregates, a collective motility of cells within the aggregate and aggregate merging is also observed. In Figs. 6E-H, we analyze the merging trajectories in the simulation. In this case, the motion is also well described by a constant acceleration model. However, in contrast to the experiment, the aggregate size dependence of the acceleration is not found to be statistically significant, with a Spearman rank-order correlation coefficient of 0.01 and a p-value of 0.962. Nonetheless, the average speed and trajectory distance divided by displacement probability densities, for both merging and non-merging aggregates (Figs. 6G and H) resemble the experiment. Notably, the trajectories in the simulation for non-merging aggregates are less directed. Finally, the merging frequency in the simulation is lower than in the experiment, with only 6% of spots merging in the simulation, compared with 35% in the experiment.

Aspects involving the motility switch were investigated systematically (SI Figs. 12, 13 & 14). Variation or withdrawal of aspects of the motility mechanism do not change the spatial characteristics of the pattern (SI Figs. 12 & 13). This highlights the fact that the spatial order of the pattern is a direct consequence of chemotaxis. An analysis of the effect of motility on merging was also conducted (SI Fig. 14). Removing the motility switch altogether results in approximately twice the merging events, but merging and non-merging aggregates does not yield distinct average speed distributions. Thus, the motility switch improves the agreement between simulation and experiment. In addition, the decrease of merging events due to the motility transition prolongs the lifetime of the aggregate pattern. Finally, removing the ability of agents to random walk after regaining motility leads to approximately three times the merging events, but the non-merging aggregates are effectively immotile. This is expected, as cells are less likely to escape an aggregate after regaining motility and thus leading to less motile aggregates.

In the simulation, merging events are caused by an imbalanced flux of agents into and out of the aggregate. An example of a typical merging event is shown in Figs. 7A-C. To keep track of the cells in each aggregate in this example, Figs. 7D and E show the changes in the number of motile and immotile cells over time. When a sufficient number of cells exits the spot and its vicinity, the core of immotile cells collectively dissolves, as the chemoattractant drops below the motility threshold (Fig. 7D arrow). This aggregate consists of only motile cells, as shown in Fig. 7B, and it moves towards and joins the other aggregate.

**Figure 7.**
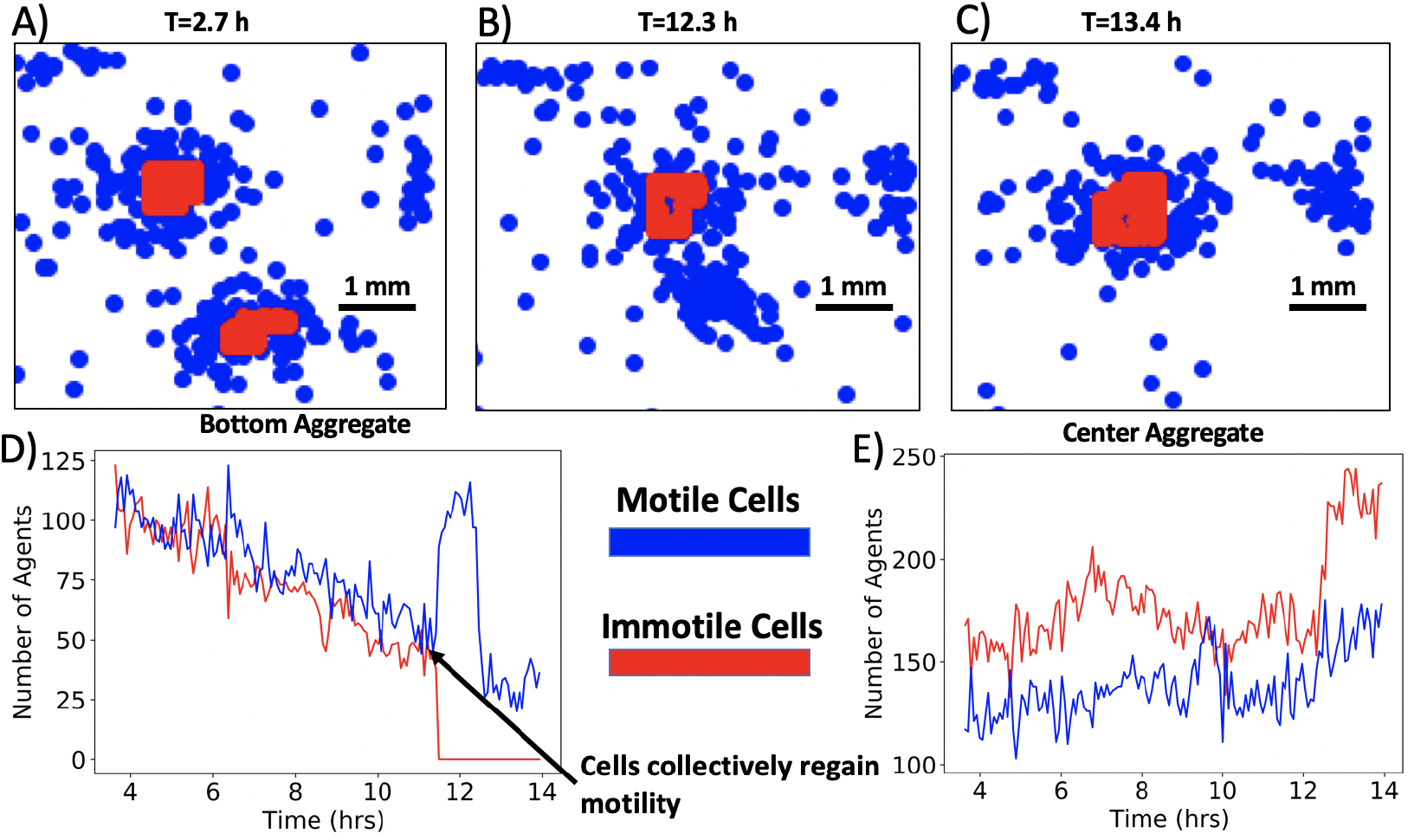
Merging at the scale of agents in the simulation. **A-C)** Three frames of the same region at different times in the simulation. The two aggregates, both consisting of motile (blue) and immotile cells (red), eventually merge. **D)** The number of cells in a 2 mm × 2 mm square region encompassing the initial position of the bottom aggregate from A) vs. time. The aggregate gradually loses both motile and immotile cells until the chemoattractant concentration drops below the threshold at 11.8 hrs (arrow). Then, all cells regain motility and collectively move towards the proximal aggregate. **E)** The number of cells in a 2 mm × 2 mm square region positioned on the initial position of the upper aggregate from A) vs. time. The aggregate experiences fluctuations in the number of cells and eventually receives the motile cells from the proximal aggregate.

In the simulations, prior to fully merging, one of the colonies becomes enriched in motile cells, which then join the other colony. This may account for the accelerated colony velocity during merging events observed in both simulations and experiments, but the experimental data does not directly show this process. Furthermore, it is consistent with the observation that merging events in the experiment lead to the formation of a new radially symmetric aggregate. The merging process depicted in Fig. 7 is ubiquitous – more examples of mergers, with visualization of the motility of the agents, are shown in SI Video 5.

## Discussion

Many examples of pattern formation and collective motility within bacterial populations have been reported, and here we examine such phenomena within populations of *Enterobacter cloacae.* This system displays two unique characteristics that have not been reported previously in the context of bacterial emergent behavior: the formation of bacterial aggregates via a transient motility transition and the collective motility of aggregates of swimming bacteria. Such behaviors are reminiscent of other bacterial systems, notably swarming and fruiting body formation in *Myxococcus xanthus.* Fruiting bodies are complex aggregates of cells that form as a result of bacterial swarming, and work over the years has shown that aggregates form as a result of contact-dependent and diffusive signaling, and that cells within the aggregate display a reduced velocity and changes in cell alignment (34–36). *Enterobacter cloacae* accomplishes similar complex behaviors using different molecular and physical mechanisms. First, motility in these experiments is the result of swimming, not surface motility associated with gliding or swarming (37). Second, aggregate formation and movement are associated with a density-dependent motility transition in which cells temporarily lose motility upon entering regions of high cell density. Because of these important differences, *Enterobacter cloacae* could serve as an important new model system for studying bacterial collective behavior.

The observed motility transition that occurs in regions of high cell density is potentially a unique mechanism of bacterial aggregate formation. A transition between motile and immotile cells is more typically associated with cells of multicellular organisms, such as the epithelial-mesenchymal transition (38). The active matter community has examined the cell motility and pattern formation (39–41). Cells used here demonstrated a complete loss of motility, as opposed to reduced motility in regions of high cell density, suggesting the transition is not due to physical processes such as crowding and jamming. Such a complete loss of motility has been reported in *Escherichia coli* due to depletion of oxygen (42). The loss of motility led to the formation of a ring of bacteria. Here a similar motility transition was associated with aggregate formation, and in fact the transition was shown to be transient. As shown in SI Video 4, cells that were initially immotile within an aggregate boundary regained motility as the aggregate boundary receded. Transient changes in motility have been reported for *Bacillus subtilis,* in which molecular “clock” switches immotile cells in a chaining phenotype back to a motile state after several hours (43). In that system, the timing of the transition was regulated by protein dilution via cell division. In the *Enterobacter cloacae,* it is unclear if cell division is occurring within dense aggregates, especially given the likelihood of low nutrient conditions within aggregates, so potentially another molecular mechanism would be needed to regulate such a transition.

The potential benefits of aggregate formation, especially aggregates composed of immotile cells, remain unclear. Previous studies have proposed that the clustering of cells into high density aggregates and biofilms is a stress response that enables cells to survive under harsh conditions such as low availability of nutrients or exposure to toxins (41,44). Would additional benefits be conferred to aggregates of immotile cells? Aggregate formation does not require a motility transition, although as suggested by the agent-based model developed here, the motility transition did produce a more stable pattern of aggregates. Immotile cells may be more energetically efficient. The length scale of the aggregates could also be set by adjustment of the molecular mechanisms that set the threshold density and timing of the motility switch, see SI Appendix 2. The transient nature of the observed motility switch did enable migration and merging of the aggregates once formed. The scheme for migration suggested by the experiments is that unbalanced fluxes of cells on different sides of the aggregate resulted in collective movement of the aggregate. Whether such symmetry breaking was random or the result of asymmetry in the chemical and environmental conditions surrounding the cell remain unclear. Potentially, if the direction of the movement of individual aggregates over multiple aggregate lengths is biased by chemical conditions, this may serve to position cells in a more favorable location. The merging process after formation may also be a mechanism to optimize the pattern of aggregates for maximum benefit to the cells.

## Materials and Methods

### Bacterial strains and culturing conditions

*Enterobacter cloacae Ecc1* used for the experiment was isolated from the Caltech turtle pond and confirmed using 16s rRNA analysis (6). Primary cultures of *Enterobacter cloacae Ecc1* were grown overnight in Luria-Bertani broth (Difco™) at 37°C under constant shaking at 180 rpm. Cultures were then washed thrice with 1X PBS (VWR, life science) and resuspended in 5 ml 1X PBS before inoculating in M9 minimal agar (6).

### Aggregate formation in BioAssay plate

To observe aggregate formation, soft agar was prepared by mixing M9 medium (Difco™) supplemented with 4% glucose (aMResco^®^) with 0.26% of bacteriological agar (Sigma-aldrich). Primary culture of *Enterobacter cloacae Ecc1* was added to sterilized minimal agar to final OD of 0.1.95.6 ml of the mix was poured into a 20 × 20 cm Bioassay dish (Thermo Fisher Scientific) with a lid. The BioAssay plate was allowed to set at room temperature for 1 hour before sealing it with parafilm.

The plate was incubated at room temperature for 2 days on a glass table. Reflection of the base of the Bioassay plate using a mirror was captured automatically by camera (Canon, EOS REBEL) every 15 minutes.

### Aggregate formation in Lab-Tek chamber

Culture was made visible by mixing *Enterobacter cloacae Ecc1* expressing RFP with *Enterobacter cloacae Ecc1* to a final concentration of 5%. Culture was mixed with 10 ml soft minimal agar to final OD of 0.0002. 2 ml of mix was allowed to set in the Lab-Tek chamber (Nunc™, Thermo Fisher Scientific) for 40 minutes, before sealing the chamber with parafilm. The entire assembly was incubated at room temperature for 24 hours, at that time visible aggregates had formed. The setup was mounted on a microscope for imaging.

Aggregates were imaged in phase contrast and RFP channels using 100X oil objective (1.5 NA CFI plan apochromat) on Nikon Eclipse Ti-E microscope with a sCMOS Camera (Zyla 5.5 sCMOS, Andor) at a 1 minute time interval with an exposure time of 30 and 100 ms respectively.

### Measurement of growth rate

To measure the growth rate of *E. cloacae Ecc1* transformed with pZE25-RFP and wild-type *E. cloacae Ecc1,* primary cultures of both the strains were inoculated in 100 ml Luria-Bertani broth at 1% inoculum in triplicate. The cultures were incubated at 37°C at 180 rpm. Absorbance of the cultures were measured at 600 nm using colorimeter (Spectronics 200, Thermo Scientific) at the interval of 30 mins.

### Estimation of cellular motility

Primary cultures of *E. cloacae Ecc1* wild-type as well as transformed with pZE25-RFP were diluted ten times. 5 μl of diluted culture was spotted on a microscope slide, which was then covered with glass coverslip (22 mm × 22 mm). Movement of an individual cell in a drop was recorded using 40X objective on Nikon Eclipse Ti-E microscope with a sCMOS Camera (Zyla 5.5 sCMOS, Andor) with a time interval of 0.1 sec.

### Image analysis

The 1X plate frames were processed with background subtraction and enhanced contrast such that only 0.01 percent of pixels are saturated using imageJ (45,46). The processed frames were segmented using Weka, a machine learning algorithm for microscopic pixel segmentation (47), example segmentation shown from frame T=22.5h in SI Fig 16. Finally, dim segmented aggregates were filtered according to maximum pixel brightness, a process depicted in detail in SI Video 6. Plotting the distribution of maximum pixel intensity, the threshold value of 0.45 was chosen to remove detected regions that were not high density aggregates.

To perform the motility and merging analysis, tracking analysis was performed in imageJ via the Trackmate plugin (48). The software is capable of producing tracks for all the segmented aggregates. Trajectories that did not lead to a merger were analyzed to determine the motility of non-merging spots. Trajectories that only contained merging events with strictly two aggregates were subject to the merging analysis. Although the aggregate size was also identified by Trackmate, the aggregate size as determined from the segmentation was used instead for greater precision and accuracy.

Images acquired at 100X were analyzed to evaluate the speed of aggregate movement using ImageJ 1.53a (45). Initial and final time frames were Otsu thresholded to find aggregate boundary, which was then marked using a freehand line tool. XY-coordinates of the centroid of the resultant line were extracted and used to calculate the distance covered by the progressing aggregate front using the Euclidean distance formula.

To measure motility of an individual cell of *E. cloacae Ecc1,* time-lapse movies of bacterial movement were opened in ImageJ 1.53a. The centroid of a single cell was tracked manually over time and its XY-coordinates were used to calculate temporal displacement of the cell, which in turn was used to measure the velocity of the cell per second.

### Radial pair correlation and structure factor

The pair correlation function also known as the radial distribution function was calculated by considering the point patterns generated by the aggregate positions. All aggregates were considered in a given square region, except for aggregates that lie within a radius of *r_max_* from the boundary. For each point, the distance between all other points was calculated. Then, the unnormalized probability density distribution was obtained for all the distances obtained. To obtain the probability density, the bins were set according to the desired resolution, defined from 0 up to and including *r_max_* in increments of *dr*. Furthermore, the probability density was divided with the number density of the pattern, *ρ* = *N_total_*/*L*^2^, where *N_total_* is the total number of points and L is the length of the region. Finally, the value in the nth bin was divided with the area of the respective annulus that spans from the bin range, with an inner radius of *r_in_* = *n * dr* and an outer radius of *r_out_* = (*n* + 1) * *dr*. Finally, the values of the resulting probability density was averaged for all points to retrieve the pair correlation function. Furthermore, the structure factor was obtained by taking the fourier transform of the correlation. Thus, by integrating the pair correlation function we can obtain the structure factor: 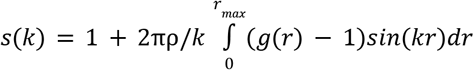(49).

### Simulation

The simulation code was written in Cuda, compiled in gcc version 4.9.4 and run on Cuda version 9.2.88, executed on a single GPU node. System size was set to a 384×384 pixel grid with periodic boundary conditions. At the beginning of the simulation, 100,000 motile agents were placed on random points in the grid. The chemoattractant concentration was initialized to a value of zero. The simulation final time was set to 2000 with a step size of Δt=0.005. Chemoattractant concentration, agent position and velocity were outputted in text file format for further analysis and plotting. The main parameter set used can be found in SI Table 1, and is termed the control parameter set.

The output of the simulation was segmented and tracked. To segment the aggregates from the simulation results, a Matlab script was written. For each pixel, the number of agents within a radius of 3 pixels was determined. When the number of agents exceeded 40, the pixel was considered to belong to an aggregate, an example segmentation is shown in SI Fig 17. In addition, aggregates that consisted of less than 10 pixels were discarded. To track the aggregates, the segmented images were entered into imageJ where the Trackmate (48) plugin was used to track the aggregates and output the trajectories.

## Acknowledgements

We would like to thank Marcel Meyer for helpful discussions and sharing the computational framework. Computation for the work described in this paper was supported by the University of Southern California’s Center for High-Performance Computing (carc.usc.edu). Osman Kahraman was also helpful in getting this project off the ground. JB acknowledges support from NSF CAREER Award PHY-1753268.

## Supplemental Information

**SI Video 1. Whole plate timelapse.**

**SI Video 2. Experimental replicates.** Video shows spot formation in three different set ups under identical growth conditions at the time interval of 15 mins. A few merging aggregates in three different plates have been indicated with cyan arrows. The middle video shows the plate that was used for all analysis shown in paper. Scale bar-10 mm. Time stamp-hrs: mins.

**SI Video 3. Aggregate advancing edge.** Microscopic images of aggregate movement captured in RFP channel with 1 min time interval. Solid yellow line indicates the progression of the aggregate front. Scale bar-40 μm. Time stamp-hrs: mins.

**SI Video 4. Aggregate receding edge.** Microscopic images of the aggregate movement recorded in phase contrast channel with frames 1 min apart. Trailing front of the spot has approximately been denoted by the solid yellow line. Scale bar-40 μm. Time stamp-hrs: mins.

**SI Video 5. Motility and merging.** In the agent based simulations, the process that leads to merging in simulations is ubiquitously described by a collective regain of motility of the aggregates. In the video, multiple merging events take place. Cells marked in red are immotile and in blue are motile.

**SI Video 6. Thresholding dimmer aggregates.** In this video of a zoomed in region of the plate, the processing step of thresholding dimmer aggregates is shown for the course of the experiment. The left panel highlights aggregates that were kept for further analysis with blue and aggregates that were not considered with red. The right panel shows the same image as the left, but without highlighting spot boundaries, for a clearer view of the aggregates.

**SI Appendix 1. Mathematical description and simulation components**

**SI Appendix 2. A model for steady state colony size**

**SI Table 1. Parameters for simulation**

**SI Figures. Supplemental figures.**

## Supplemental Figures

**SI Fig 1.**
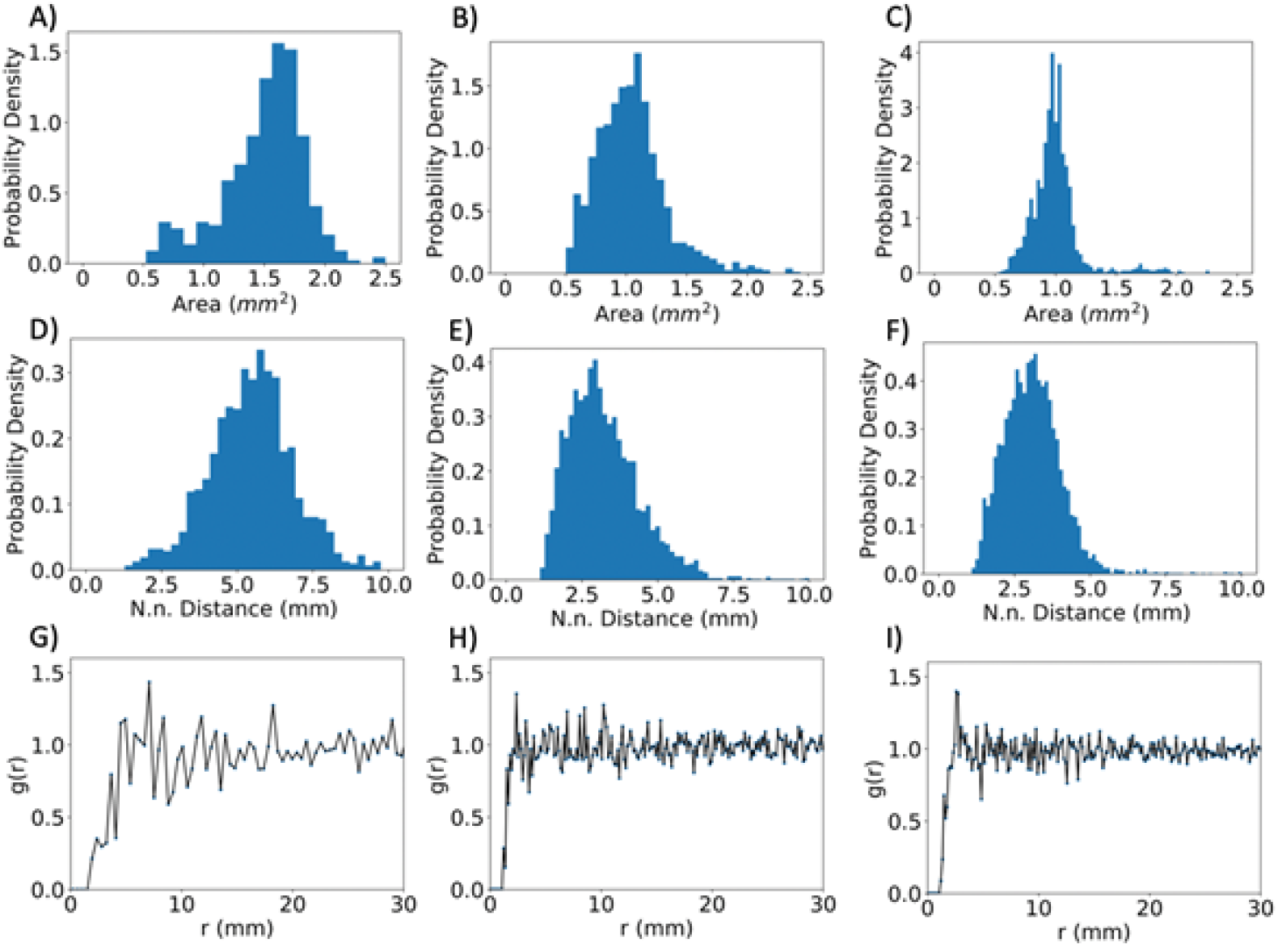
Distribution of areas, nearest neighbors and pair correlation functions of aggregate positions for replicate experiments. Spatial quantities are calculated for three different plates at 22.5 hrs. The first two columns are unused replicate experiments and the third column is obtained from the plate used throughout the study. The quantities are calculated for a 10×10cm subregion in the center of the plate to account for edge effects. **A-C)** Aggregate area distributions. **D-F)** Nearest neighbor distance distributions. **G-H)** Pair correlation functions.

**SI Fig 2.**
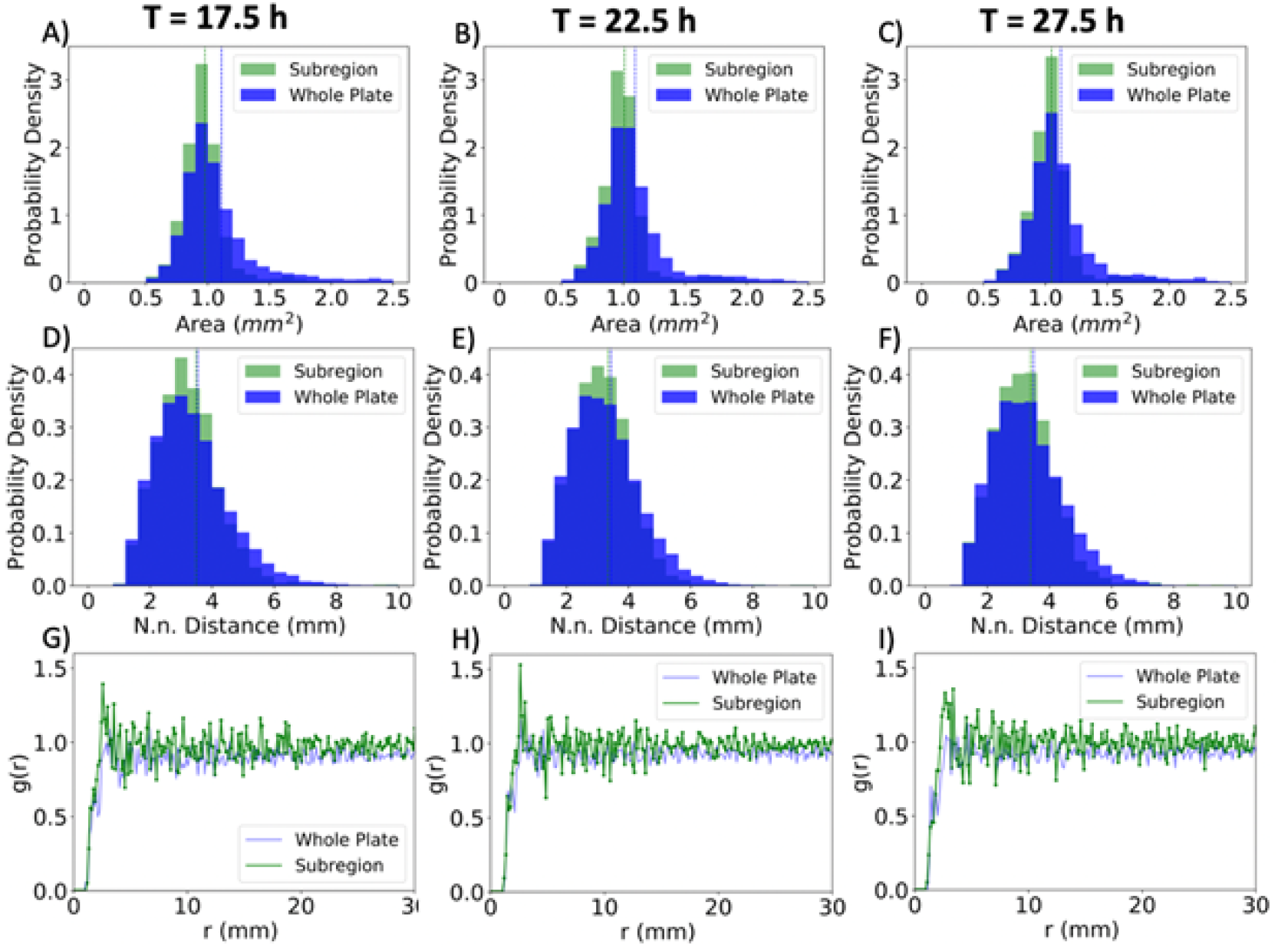
Distribution of areas, nearest neighbors and pair correlation functions for aggregate positions at different times for both the whole experimental plate and the subregion of interest. Spatial quantities for three time points 17.5 h, 22.5 h and 27.5 h from left to right. The quantities were plotted with blue for the whole plate and green for the subregion used in the main text, shown in SI Fig 2. **A-C)** Aggregate area distributions. **D-F)** Nearest neighbor distance distributions. **G-H)** Pair correlation functions.

**SI Fig 3.**
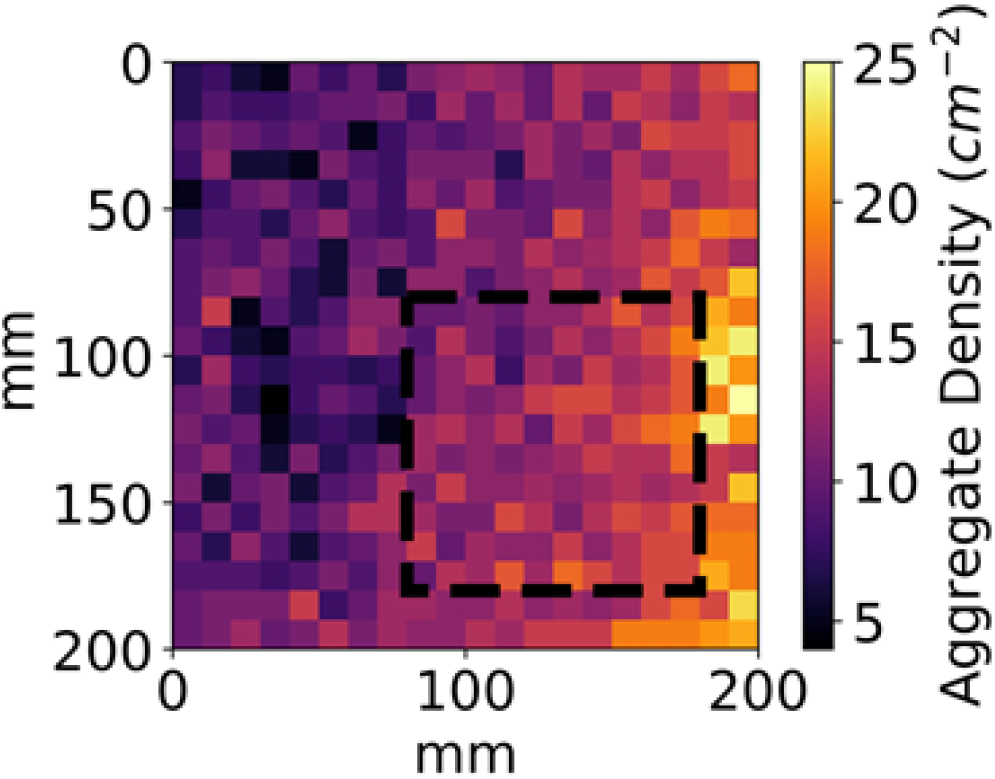
Spatial aggregate density fluctuations. The aggregate number variations are visualized for the frame used for the spatial analysis, at T=22.5 h. The highlighted region marks the spatial subset of the plate analyzed in Figure 2.

**SI Fig 4.**
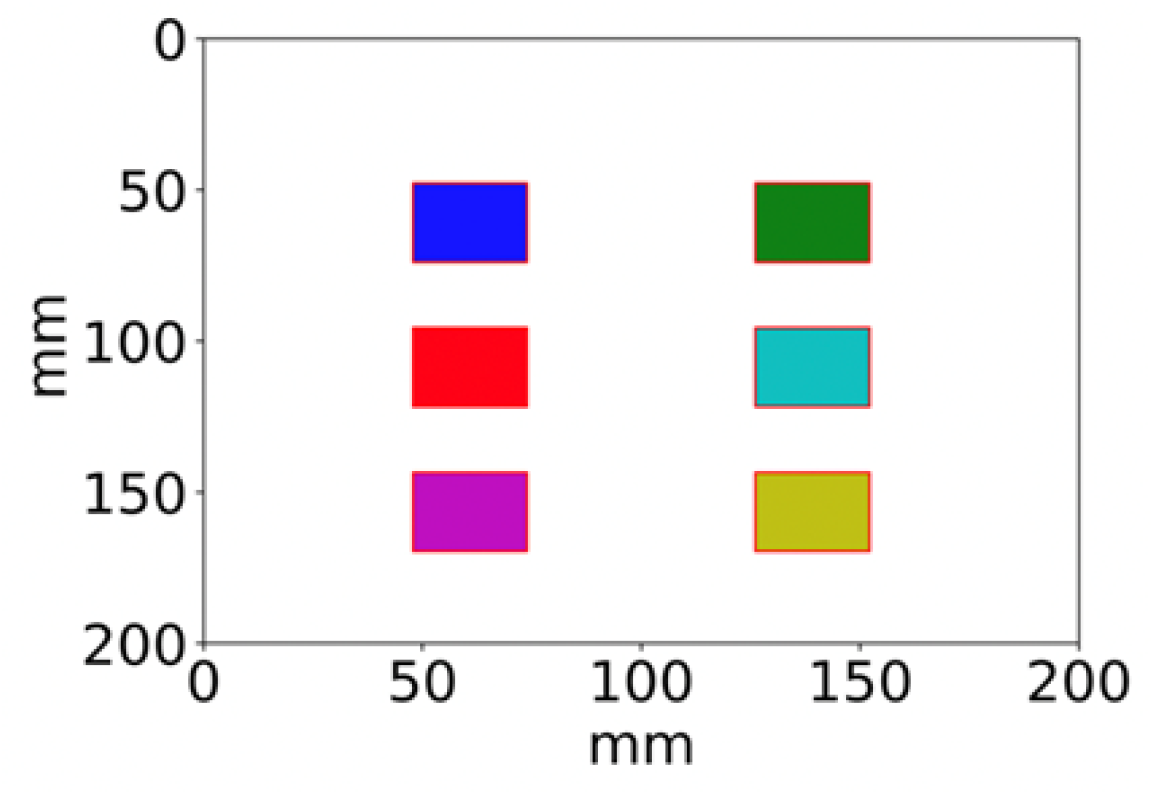
Regions of the plate used for analysis of motility and merging.

**SI Fig 5.**
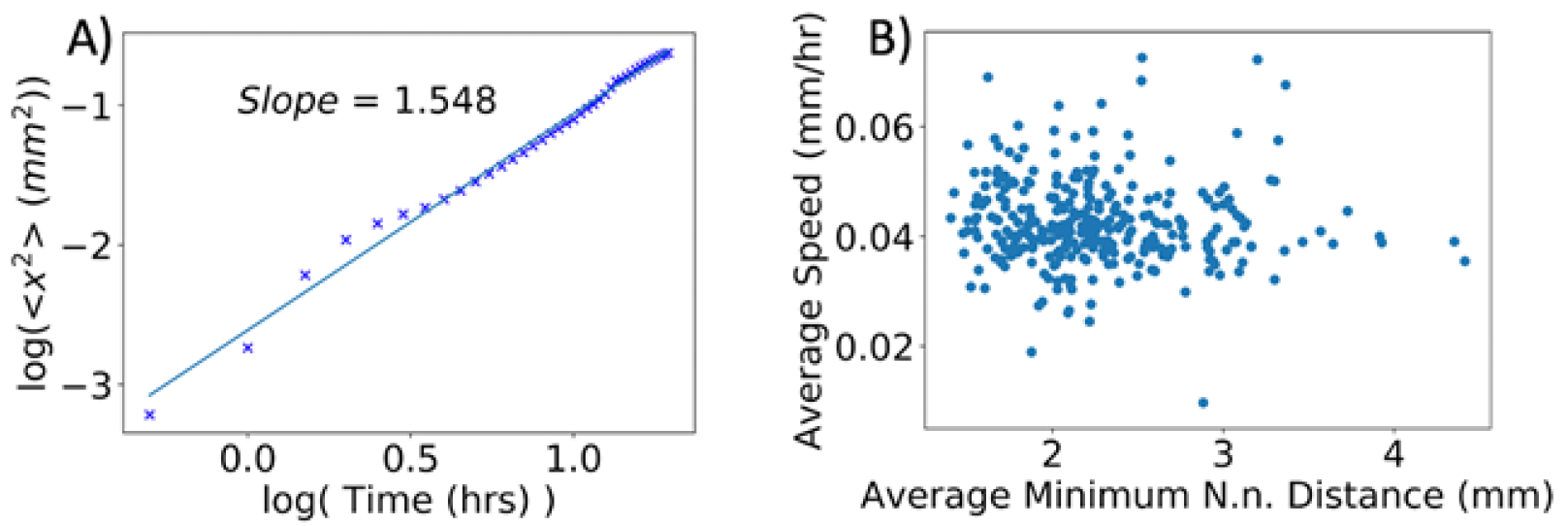
Trajectory characterization. **A)** Log-log plot of mean square displacement versus time. The fit yields a coefficient of 1.5 indicating that the trajectories are, on average, superdiffusive. Brownian motion, corresponding to diffusion, yields a coefficient of 1.0. Thus, the aggregate trajectories are directed. **B)** Average speed versus average minimum nearest neighbor distance for all analyzed aggregate trajectories. To obtain the latter quantity for a single trajectory, the nearest neighbor distances were calculated for each frame, the minimum was extracted and averaged over all frames. Spearman rank-order correlation coefficient correlation yielded −0.12, with a p-value of 0.034.

**SI Fig 6.**
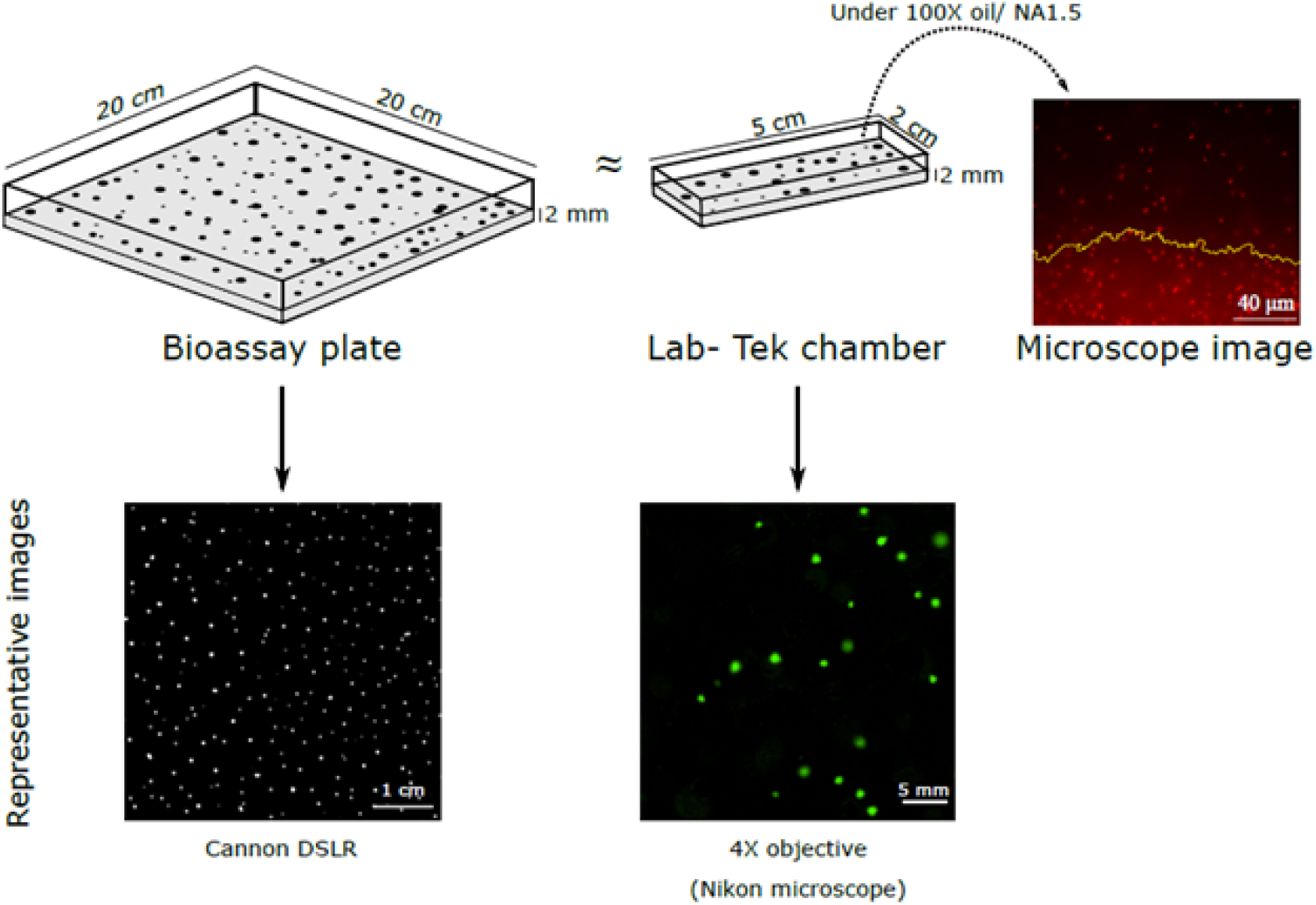
Depiction of experimental set-up using bioassay plate and Lab-Tek chamber. The height of agar in both the vessels was 2 mm. Representative images taken by using DLSR camera and 4X objective on Nikon microscope for respective set-up has been shown. Spot formation in Lab-Tek chamber was imaged in time-lapse microscopy using 100X oil/ 1.5 NA objective.

**SI Fig 7.**
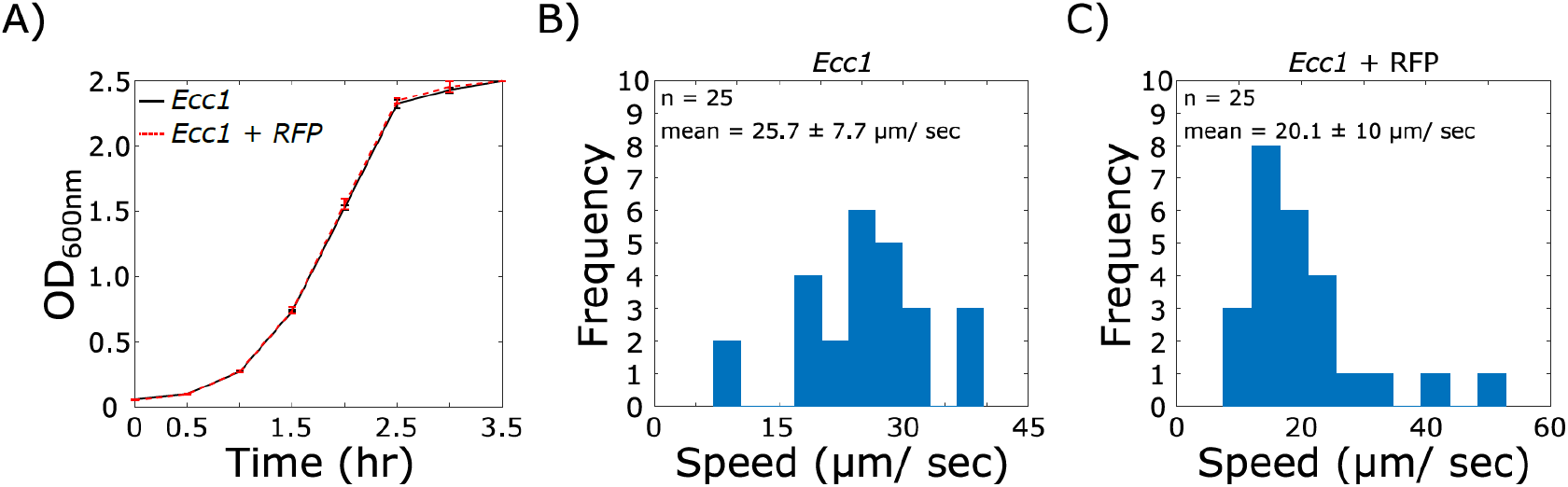
Control measurements for RFP producing populations. *Enterobacter cloacae* Ecc1 cells transformed to express RFP protein were compared with *Enterobacter cloacae* Ecc1 cells for growth rate (A) and single cell motility (B and C).

**SI Fig 8.**
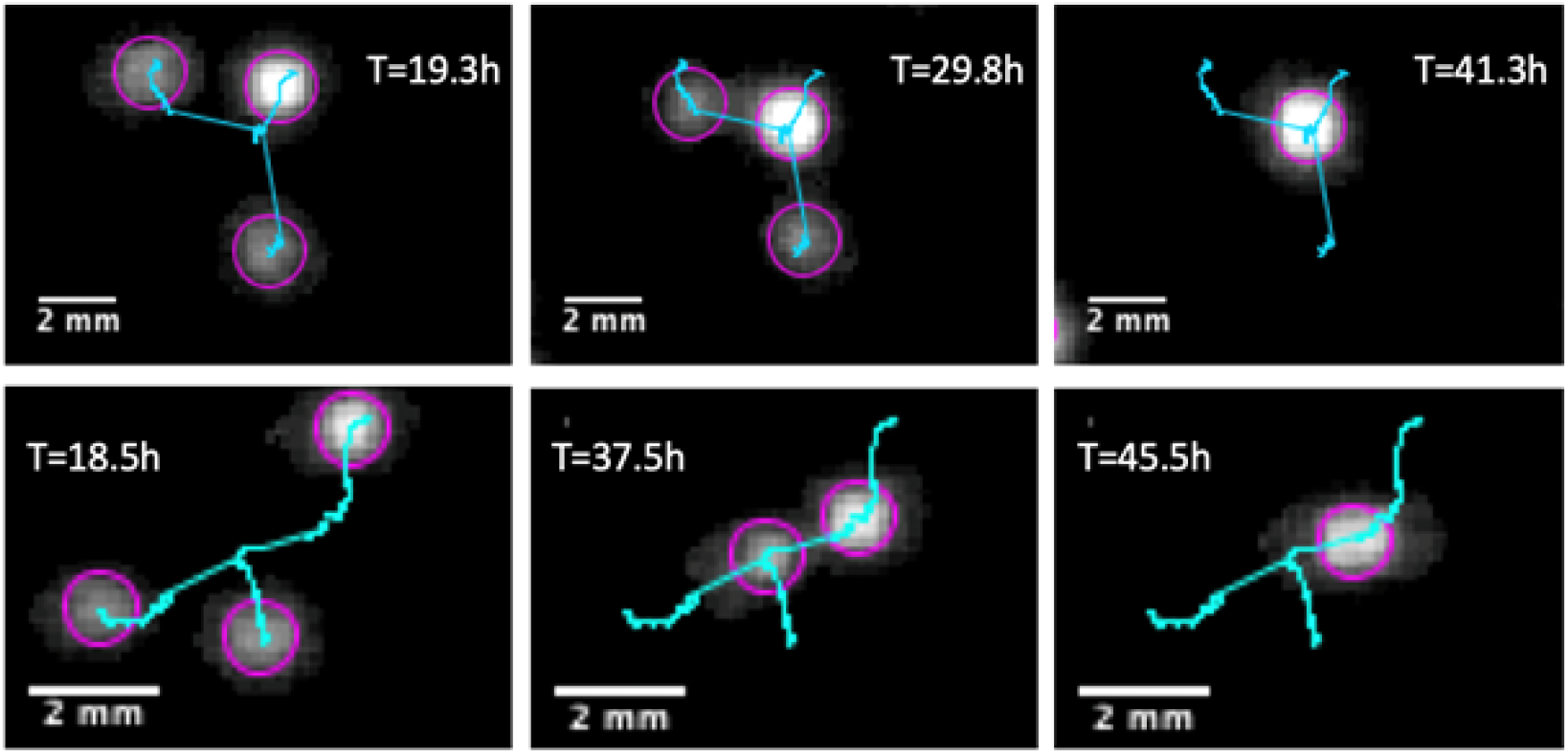
Three aggregate merging instances. Of all the aggregates tracked, only 4.2% took part in a merger involving three aggregates. Two instances of a three aggregate merger are depicted at three characteristic time points.

**SI Fig 9.**
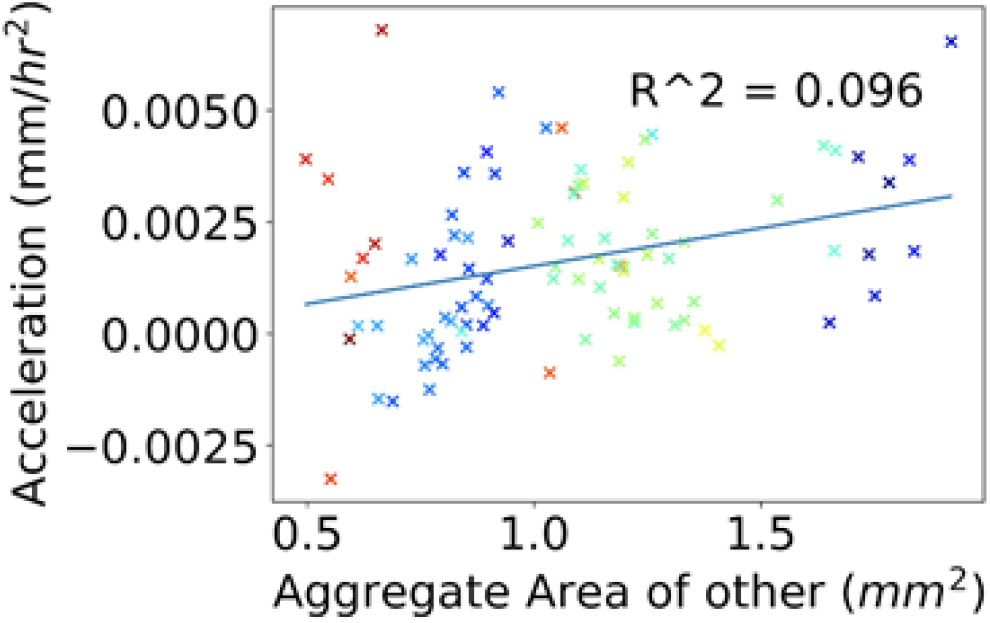
Aggregate acceleration during merging when analyzed with respect to collision point. This is an additional analysis performed on two-spot merging trajectories observed in experimental measurements. In this instance, the distance of each spot relative to the collision point versus time is fitted to a quadratic equation. Approximating the trajectory as constant acceleration motion, the acceleration is obtained from the quadratic equation. Finally, it is plotted with respect to the other merging aggregate area. The two quantities have a spearman rank-order correlation coefficient of 0.31 with a p-value of 0.002. The color of each data point corresponds to a merging event taking place in the respectively colored subregion in SI Fig 4.

**SI Fig 10.**
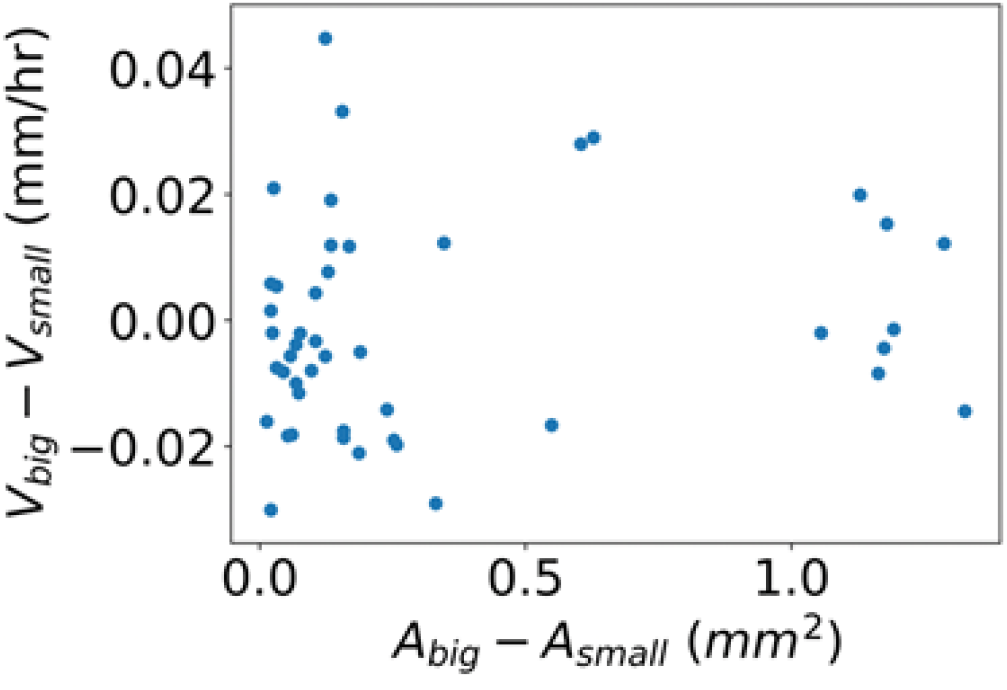
In this plot of experimentally derived quantities, for each two-spot merging event, the difference in average speed is plotted versus the difference of aggregate size between the two aggregates. Small aggregates do not necessarily move faster than big aggregates during two-spot merging events. This is the case since the difference of speeds is not always positive, even when the size difference is significant.

**SI Fig 11.**
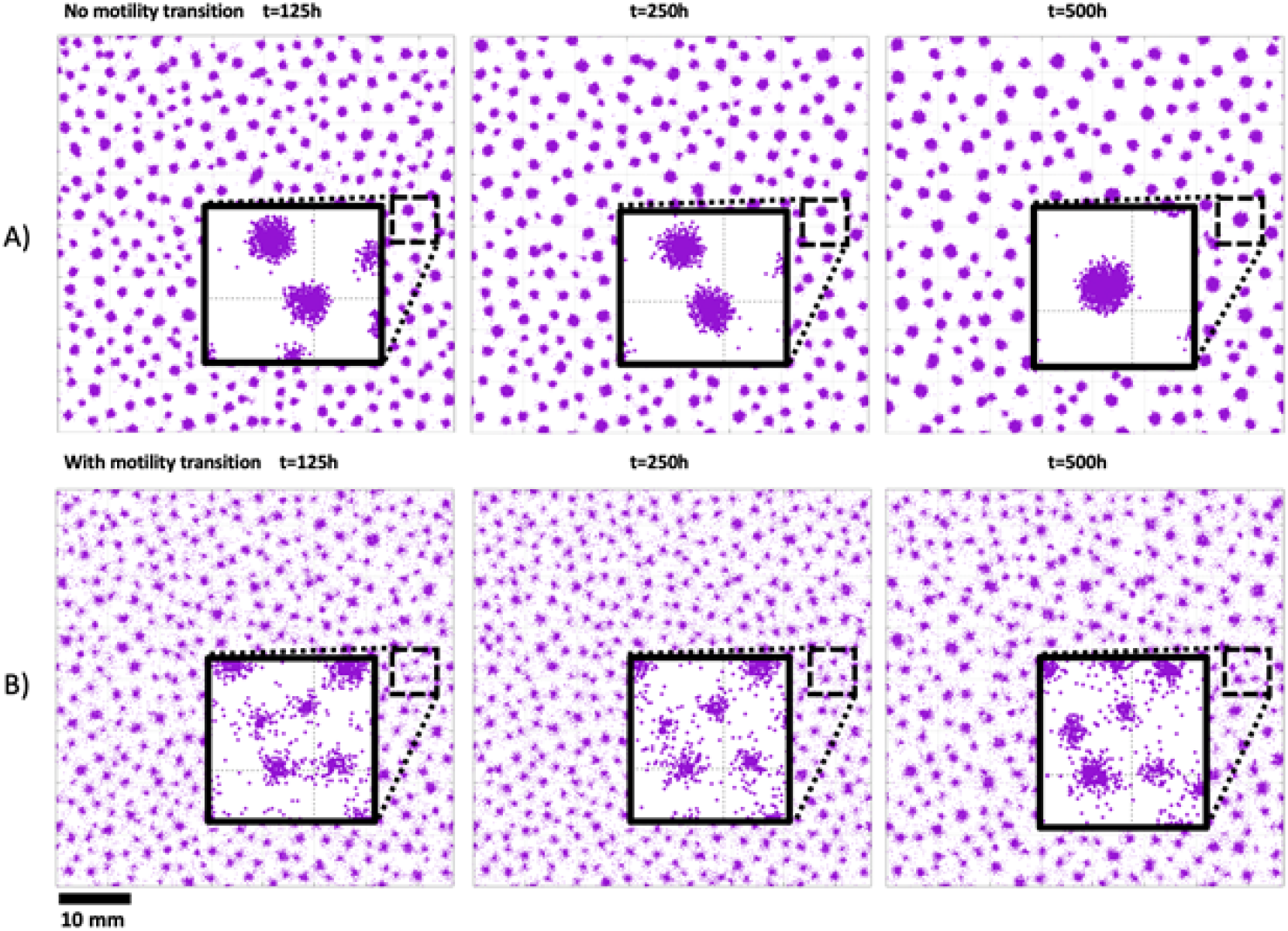
In the simulation, introducing a motility transition dramatically increased the proportion of cells not bound in aggregates over long time scales. **A)** With no motility transition. **B)** With the motility transition.

**SI Fig 12.**
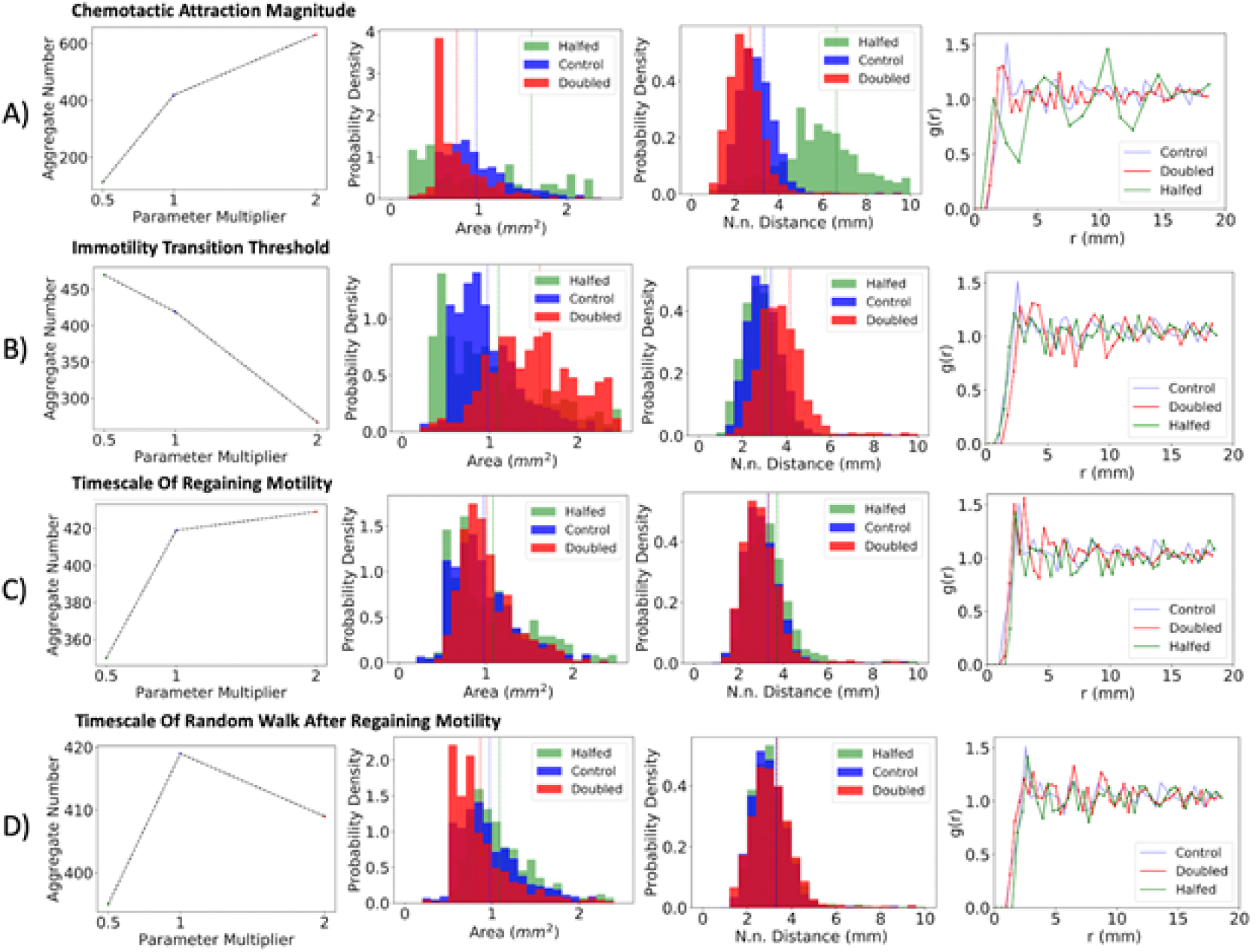
Effect of varying essential parameters of the simulation on spatial structure. **A)** Chemotactic attraction magnitude **B)** Immotility transition threshold **C)** Timescale of regaining motility **D)** Timescale of random walk after regaining motility.

**SI Fig 13.**
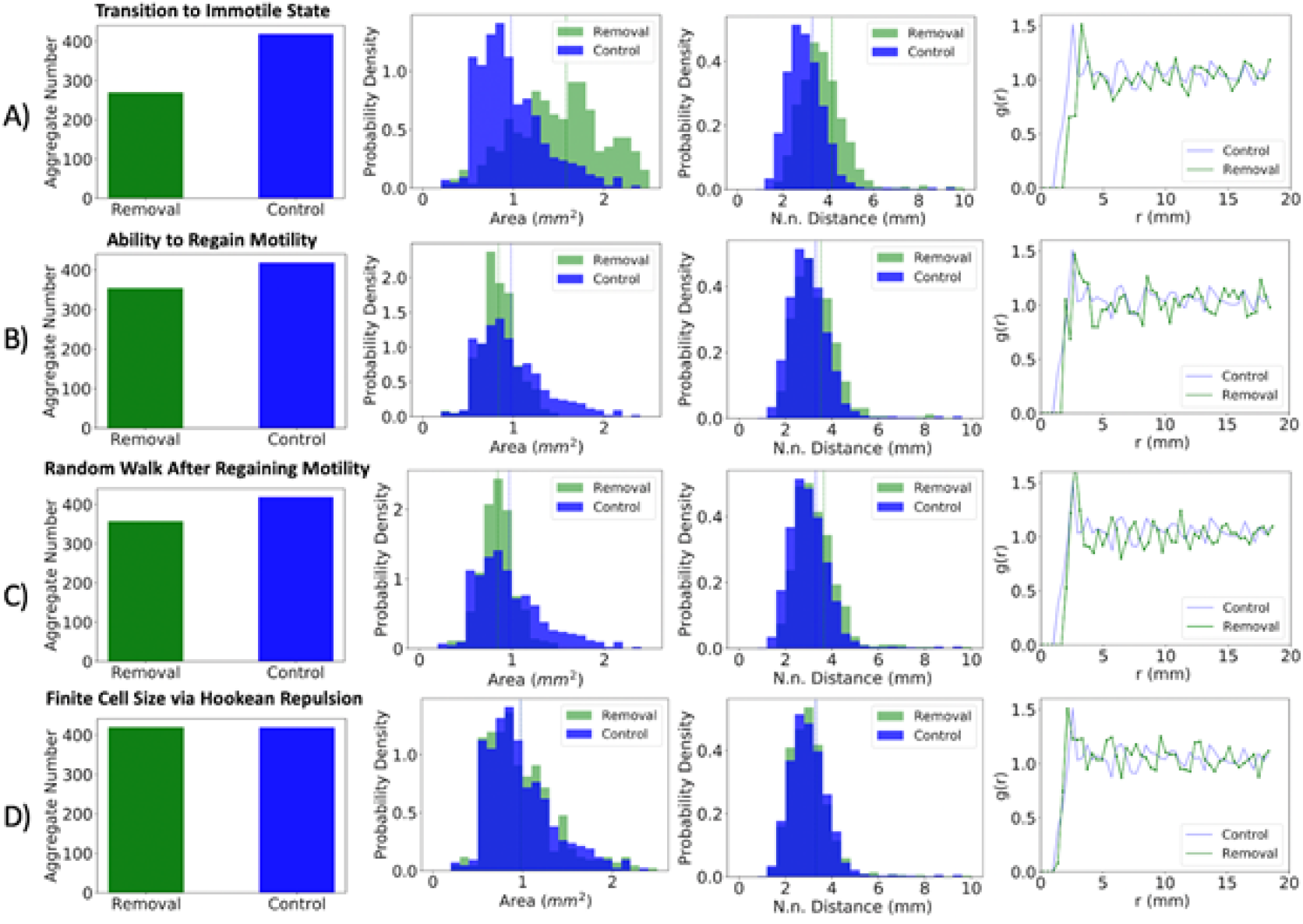
Effect of removing aspects of the simulation on spatial structure. **A)** Chemotactic attraction magnitude **B)** Immotility transition threshold **C)** Timescale of regaining motility **D)** Timescale of random walk after regaining motility.

**SI Fig 14.**
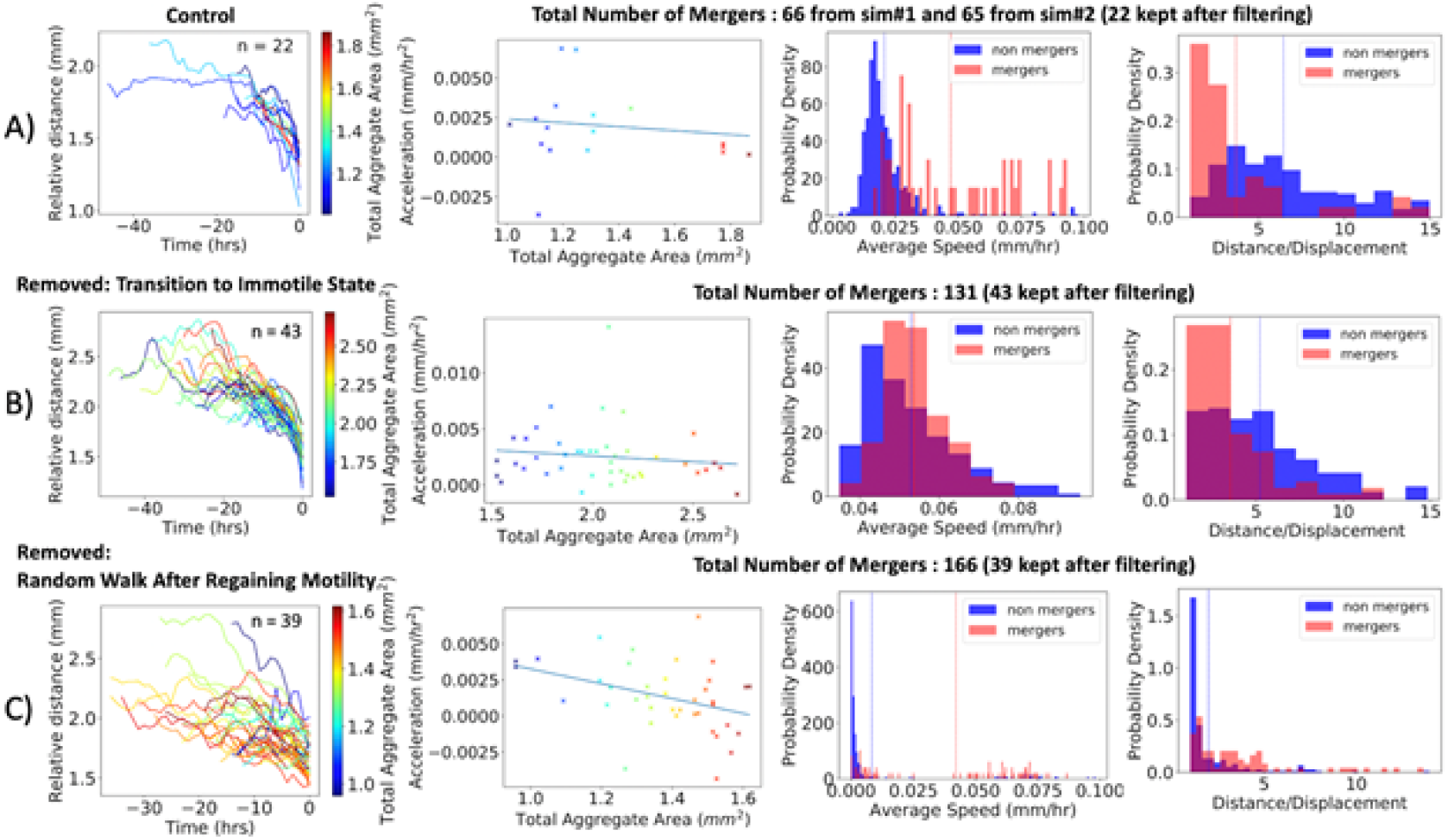
Effect of removing aspects of the simulation on motility and merging. **A)** Original simulation results for comparison **B)** Removing the introduced motility transition **C)** Removing random walk process after cells regain motility.

**SI Fig 15.**
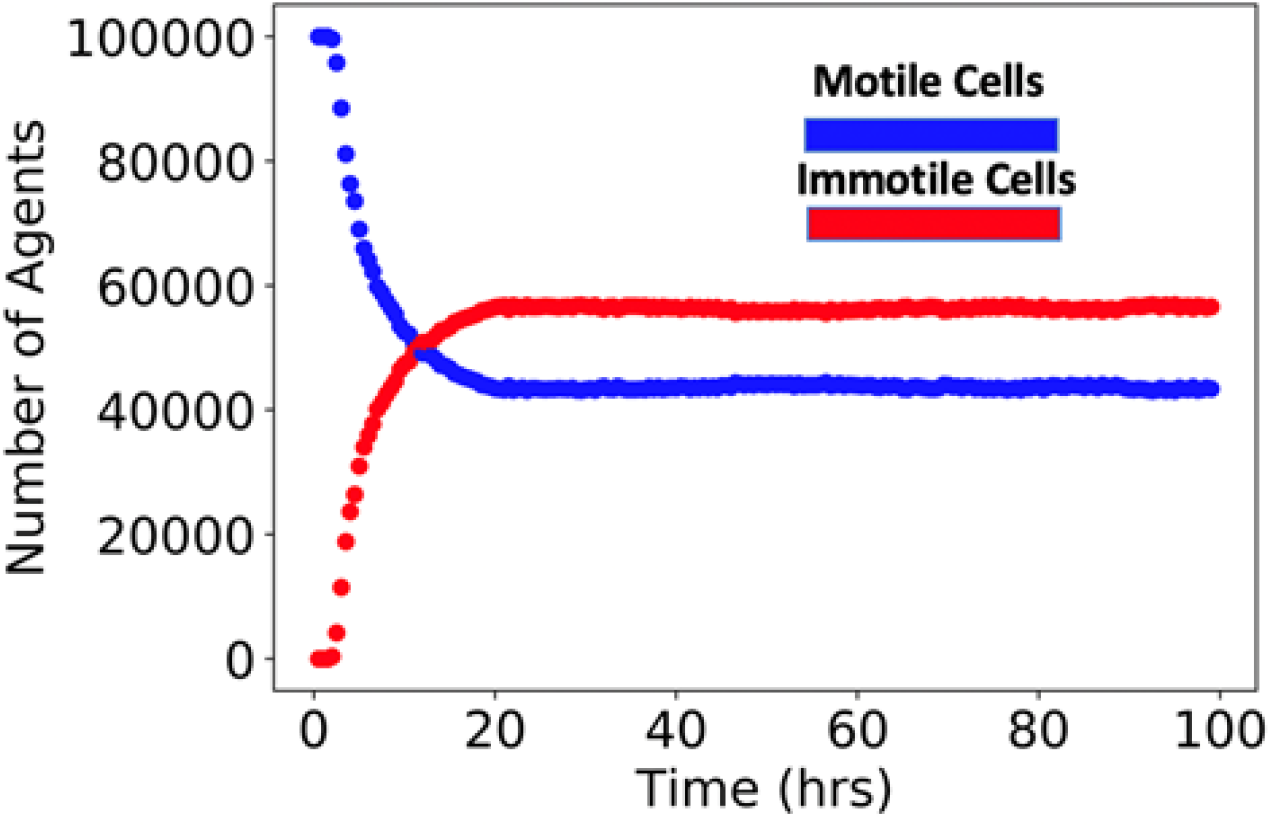
Evolution of motile and immotile cells in the simulations. During the simulations, the distribution of motile and immotile cells reaches and retains a mixed steady state.

**SI Fig 16.**
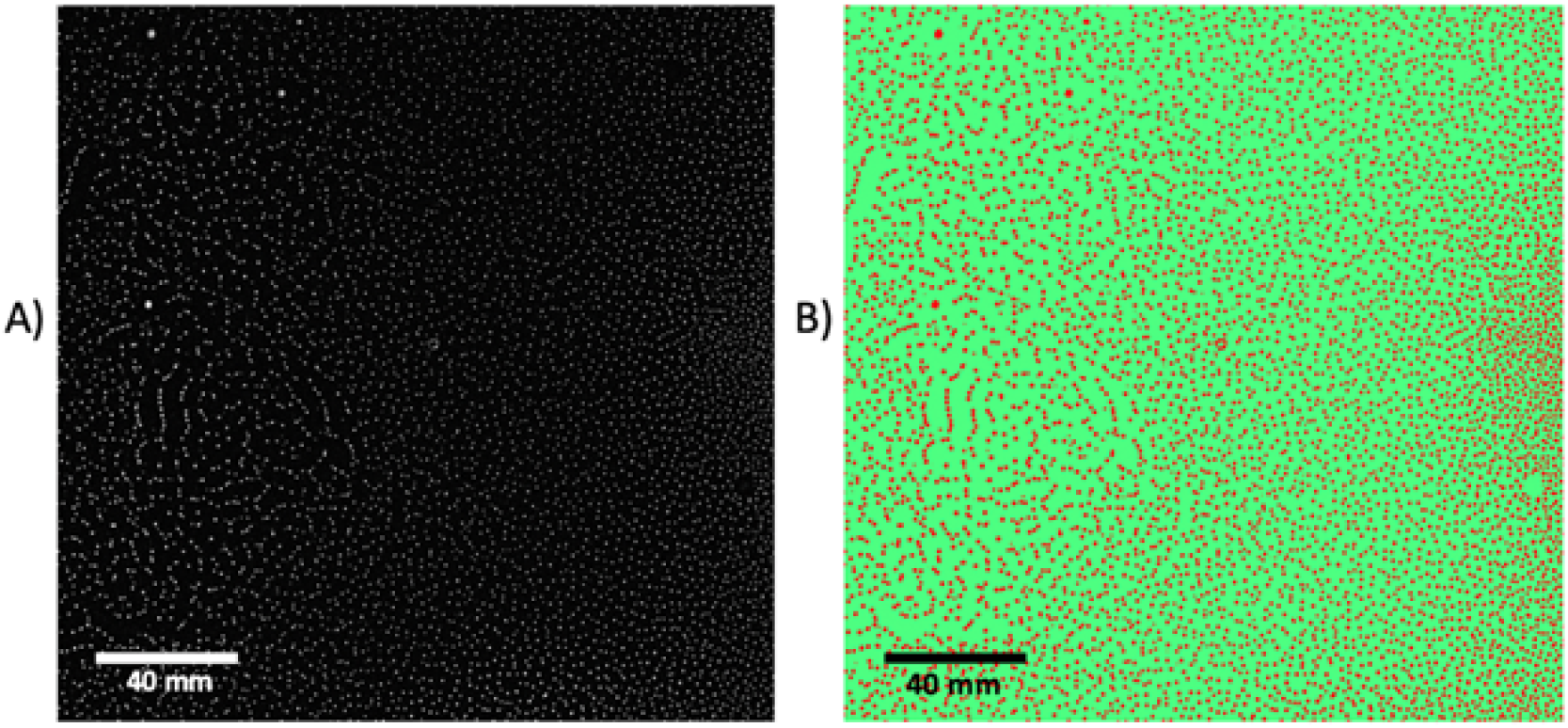
Identification of aggregates using Weka segmentation. A) Experimental image of the entire plate at T = 22.5 h. B) Aggregates identified after using Weka software.

**SI Fig 17.**
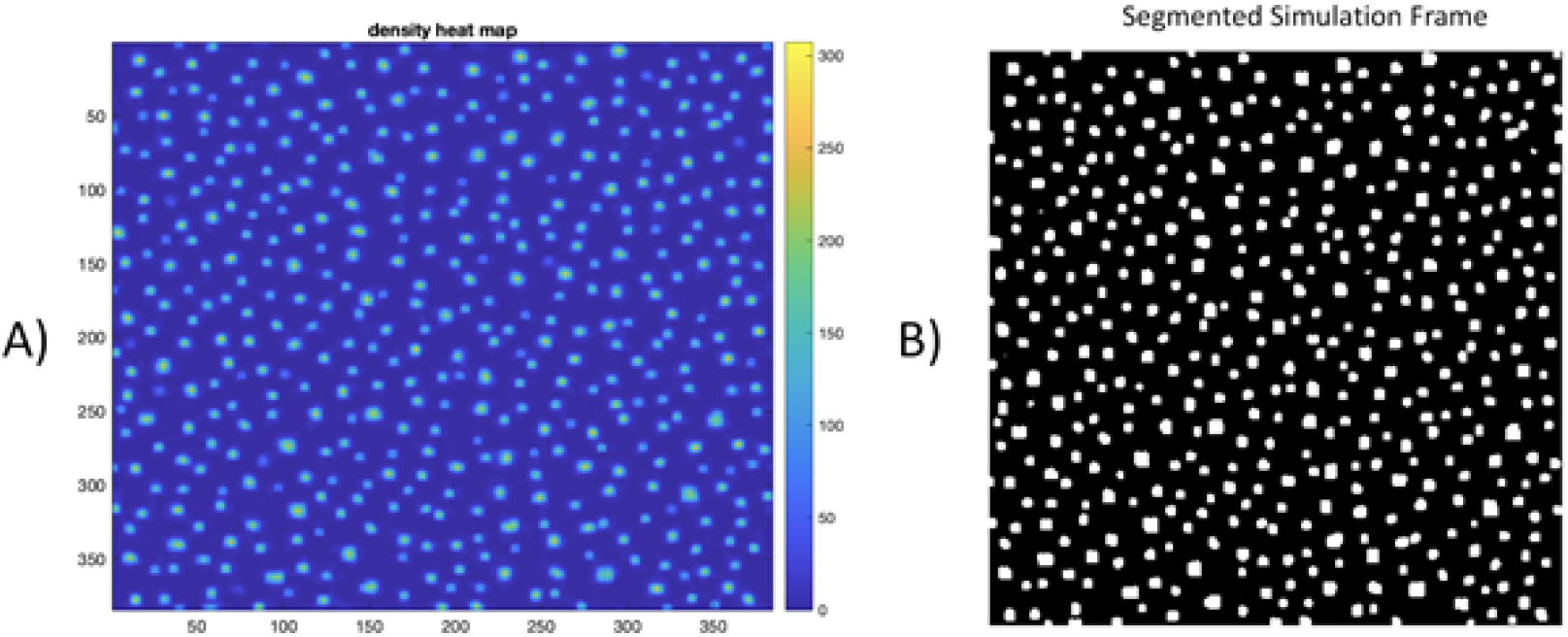
Segmentation of the simulation. A) Density heat map of simulation results. For every pixel, the number of agents within a radius of 3 pixels is used to calculate the local cell density. B) Segmentation results after only considering pixels with a density value greater than 40 and aggregates with an area greater than 10 pixels.

